# Instantly Reversible Enzyme Regulation via Dichromatic Irradiation of Incorporated Azobenzene

**DOI:** 10.1101/2024.07.04.602025

**Authors:** Ranit Lahmy, Caroline Hiefinger, Fjoralba Zeqiri, Sabrina Mandl, Willibald Stockerl, Ruth M. Gschwind, Burkhard König, Andrea Hupfeld

## Abstract

Azobenzene is a widely recognized tool for achieving artificial spatiotemporal control of enzyme activity using light. Photocontrol reversibility is typically based on photostationary states with varying *E* and *Z* isomer compositions achieved through irradiation at specific wavelengths. Here, we report an alternative mechanism for azobenzene based enzyme regulation, discovered through simultaneous irradiation with two wavelengths. Using two engineered variants of imidazole glycerol phosphate synthase, in which azobenzene was incorporated as an unnatural amino acid to enable reversible control under monochromatic irradiation, we uncovered unique behavior under dichromatic irradiation. Notably, a distinct spectroscopic signal from the azobenzene moiety emerged during simultaneous irradiation at 365 nm and 420 nm and vanished upon return to the dark. Intriguingly, dichromatic irradiation triggered a reproducible two-fold increase in catalytic activity and an instantaneous return to baseline activity in the dark for one variant. In contrast, the other variant and the wild-type enzyme maintained their baseline activity under the same conditions. These findings reveal an unexplored avenue for azobenzene photoswitching, offering a novel approach to photocontrol with potential applications in sequentially regulating multiple enzymes, especially when combined with monochromatic irradiation strategies.

## Introduction

The reversible regulation of enzyme activity with light is an emerging subdiscipline in chemical biology with significance for various applications including cell studies and multi-enzyme cascade biocatalysis. To establish reversible photocontrol, synthetic photoswitches are commonly integrated within ligands that are (non)covalently attached to the enzyme, or incorporated directly into the enzyme scaffold as unnatural amino acids (UAAs).^[1]^

Azobenzenes are one of the most widespread photoswitc**h**es (**Figure 1A**). They exist in a planar, thermally stable *E* isomer with C_2h_ symmetry,^[2]^ which allows a _π→π_* transition with a strong light absorbance at ∼320 nm and prohibits a n**→**_π_* transition with only weak light-absorbance at ∼450 nm.^[3]^ The non-planar, thermally less stable *Z* isomer instead comprises C_2_ symmetry,^[4]^ which weakens the _π→π_* transition leading to a decrease and a shift of the UV band to lower wavelengths, whereas the now allowed n_→π_* transition causes a stronger signal at ∼450 nm.^[3]^ Irradiation with UV light of e.g. 365 nm leads to the establishment of a photoinduced equilibrium that favors the *Z* configuration (**Figure 1B**), since both isomers absorb light at that wavelength with different intensities () and isomerize upon excitation with different quantum yields (∼ 0.16 and ∼ 0.39).^[5]^ This equilibrium is named photostationary state (PSS). Likewise, irradiation with visible light of e.g. 420 nm results in an *E*-enriched PSS again du**e** to different absorbance intensities (<) and quantum yields (∼ 0.29 and ∼ 0.45).^[5]^ Such a light induced switch between the thermal equilibrium (TEQ), the PSS^365^ and PSS^420^ is key to the reversible photocontrol of enzyme activity.

**Figure 1.**
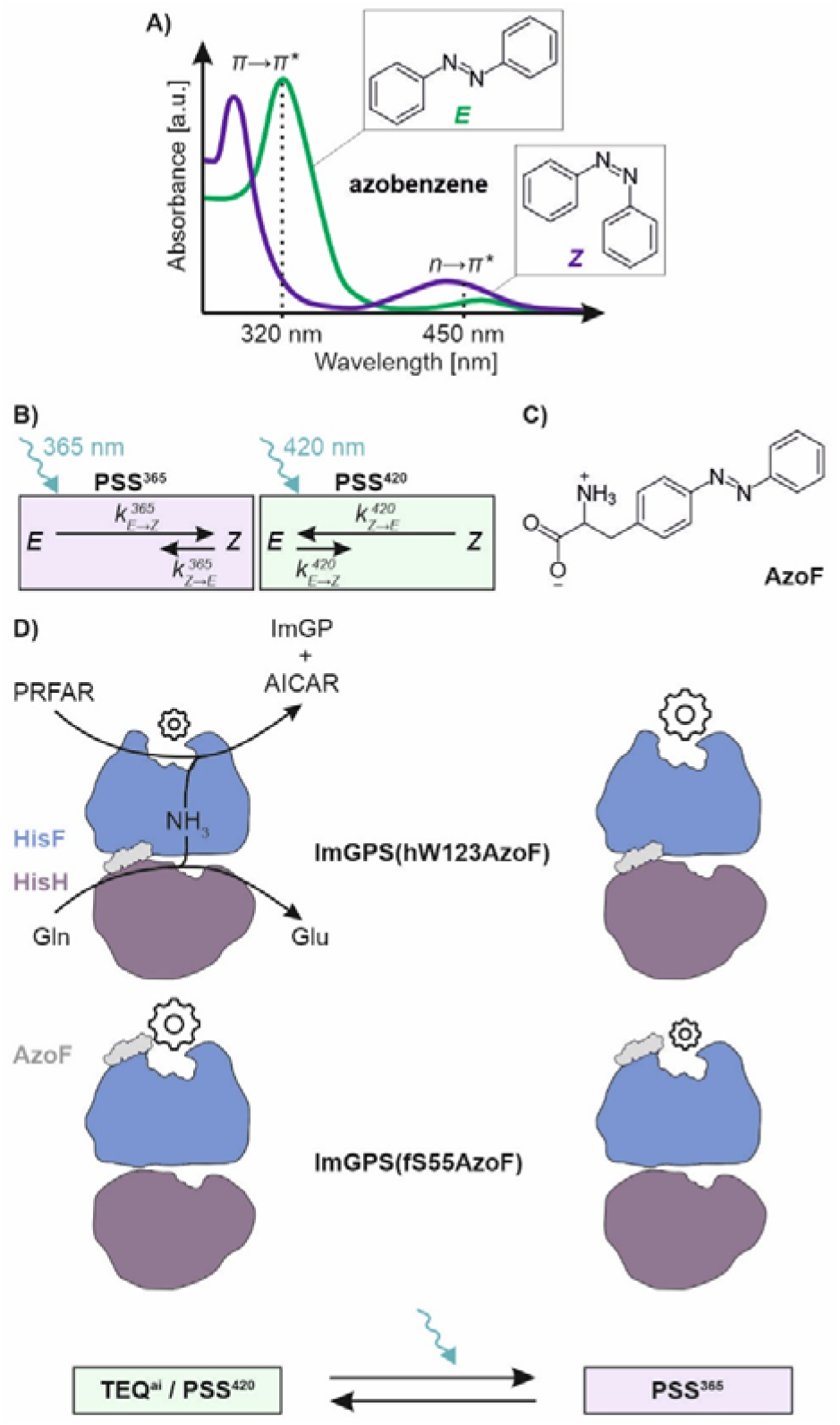
Azobenzenes and their implementation as UAA for reversible photocontrol of enzyme activity and allostery. A) Exemplary spectroscopic signal of azobenzene derivatives. B) Irradiation induces the formation of photostationary states (PSS) with wavelength specific *E*:Z compositions that depend on the isomerization rates *k* (*k* = ε *×* Φ). C) Attachment of an amino acid moiety in para position of the central C-N=N-C link generates AzoF facilitating the incorporation into enzymes. D) Subunit reactions in ImGPS and the regulation of PRFAR turnover by light in two previously engineered ImGPS-AzoF variants. Gears indicate low (small) or high (large) enzyme activity.

For example, we have recently shown that the incorporation of the azobenzene based UAA phenylalanine-4’-azobenzene (AzoF; **Figure 1C**)^[6]^ can facilitate the photocontrol of enzyme activity and allostery.^[7,8]^ Incorporation was thereby accomplished by amber suppression using a previously engineered heterologous aminoacyl-tRNA-synthetase (aaRS) / tRNA pair, which is orthogonal in prokaryotic cells.^[6]^ To achieve our goal, we used the prominent allosteric bi-enzyme complex imidazole glycerol phosphate synthase (ImGPS). ImGPS consists of a glutaminase subunit, HisH (delineated as “h”), **a**n a cyclase subunit, HisF (delineated as “f”; **Figure 1D** and **Figure S1**).^[9]^ HisH produces ammonia, which travels through an intermolecular channel to the active site of HisF where it is required for the cleavage of *N’*-[(5’-phosphoribulosyl)formimino]-5-amino-imidazole-4-carboxamide ribonucleotide (PRFAR). Notably, binding of PRFAR^[10]^ to HisF induces an allosteric stimulation of HisH that results in a drastic increase of glutamine hydrolysis.^[8]^ Our engineering endeavors identified two variants, ImGPS(hW123AzoF) and ImGPS(fS55AzoF), that demonstrated different catalytic activities in their thermally equilibrated as-isolated state (TEQ^ai^), their PSS^365^ and their PSS^420^ (**Figure 1D**).^[7,8]^ AzoF is thereby positioned either in HisH at the subunit interface [ImGPS(hW123AzoF)] or in HisF close to the PRFAR binding site [ImGPS(fS55AzoF)]. While ImGPS(hW123AzoF) exhibited a higher activity in PSS^365^ than in TEQ^ai^ or PSS^420^, ImGPS(fS55AzoF) showed a reversed activity profile. Moreover, both photocontrol effects could be associa**t**ed with conformational changes that occur within ImGPS afte**r t**he light induced isomerization of AzoF and that are connecte**d** to allostery.

Irradiation with monochromatic light has hitherto been quite effective for the photocontrol of enzymes, exemplified, inter alia, by our studies. However, we wondered how dichromatic irradiation would change the photophysical behavior of azobenzene, particularly AzoF, and how this would affec**t th**e photocontrol of enzymes. This central mechanistic question originated from the idea of injecting energy into enzymes via azobenzene excitation, which might then stimulate conformational dynamics and hence catalysis similar to heat input. We initially contemplated that AzoF might be able to frequently switch back and forth between *E*-and *Z*-enriched states by fast pulsed dichromatic irradiation, generating e.g. kinetic energy that can be passed on to the enzyme.

As a first step, we explored the consequences of dichromatic irradiation on enzyme activity with incorporated photoswitchable AzoF. To this end, we performed extensive kinetic analyses on the two previously established systems ImGPS(hW123AzoF) and ImGPS(fS55AzoF). We directly compared monochromatic 365 nm or 420 nm irradiation with dichromatic 365/420 nm irradiation. By this, we uncovered a further modus operandi of AzoF that allows a different type of photocontrol monochromatic irradiation.

## Results and Discussion

The production of hW123AzoF and fS55AzoF has than been previously established in our group.^[7,8]^ Routine procedure**s** in the course of this work confirmed the therein described properties. Both enzymes were >90% pure, contained AzoF, kept their structural integrity and showed high thermal stability with denaturation midpoints of >85°C. To show the influence of irradiation on enzyme activity, we decided to measure the full ImGPS reaction (glutamine + PRFAR _→_ glutamate + ImGP + AICAR; cf. **Figure S1**), which can be readily detec**t**ed spectrophotometrically during irradiation by following the decay of PRFAR at 300 nm. We have also tested to record the HisH activity during irradiation. While the coupled enzymatic assay that we use endured brief irradiation phases (<1 min) in our previous studies,^[7,8]^ it was significantly hampered during constant irradiation (>1 min). This is most likely due to the light-sensitivity of the flavin containing glutamate oxidase used in this assay.

We set up our experiments to detect the change in rate caused by irradiation directly during turnover with glutamine and PRFAR in saturation (cf. **Figure 2**,**3**).^[11]^ For this, we initiated **t**he reaction by the addition of ImGPS, followed the decrease of PRFAR in the dark for ∼5 min and then started the irradiation while following PRFAR turnover until the reaction stopped. To reduce the heat input by irradiation, we used light-emitting diodes (LEDs) that were powered by pulse-width modulation (“pulsed irradiation”), by which the sample is exposed to light pulses of consistent intensity (**Figure S2**). Furthermore, we use the term light-regulation factor (LRF) that compares the activity during irradiation with the activity in the dark. Finally, we performed the experiments with technical replicates, meaning more than one measurement with the same enzyme sample, and with biological replicates, meaning measurements with separately produced enzyme samples.

**Figure 2.**
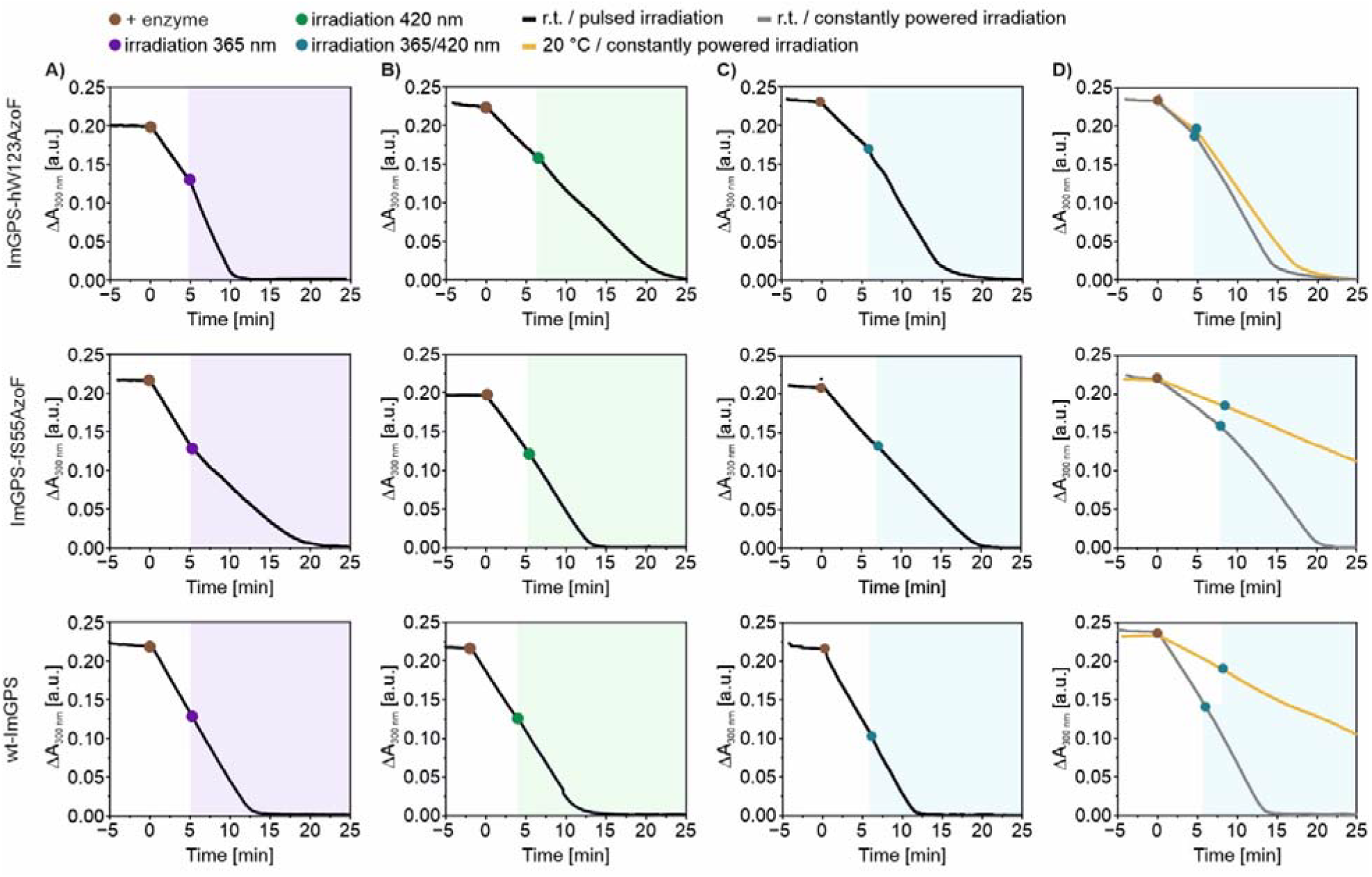
Comparison of ImGPS activity changes facilitated either by 365 nm (A), 420 nm (B), or dichromatic 365/420 nm (C, D) irradiation. For reaction conditions see **Table S1** and **Table S2**.

We started our investigation with monochromatic pulsed irradiation. The measured *k*_*cat,app*_ values (pH 7.5) for the TEQ^ai^ before irradiation were slightly higher (**Table S1**) than those reported previously (pH 8.5) owing to the pH optimum at pH ∼7.0.^[7]^ For 365 nm, we then obtained a ∼1.9-fold activity increase for ImGPS(hW123AzoF) and a ∼1.7-fold activity decrease for ImGPS(fS55AzoF), whereas wt-ImGPS activity remained unaffected (**Figure 2A**; **Table S1**). In comparison, activity values were comparable before and during irradiation with 420 nm, which establishes a similar *E*:*Z* composition as found in TEQ^ai^. (**Figure 2B; Table S1**). The observed activity increase for ImGPS(hW123AzoF) upon 365 nm irradiation clearly coincided with our previous result (LRF ∼1.9) that we acquired by irradiating the enzyme prior to the measurement with light of 365 nm.^[8]^ The same was true for ImGPS(fS55AzoF), for which we also observed an activity decrease (LRF ∼2.3) in our previous study.^[7]^

The effects observed for ImGPS(hW123AzoF) and ImGPS(fS55AzoF) during monochromatic irradiation are most likely caused by the establishment of a new *E*:*Z* composition. Since the *Z* isomer shows only slow thermal relaxation within hours to days this change in activity should be maintained when irradiation stops after a few minutes. To confirm this, we used **a**n experimental setup, in which we initiated the reaction by addition of the enzyme, followed PRFAR turnover for ∼5 min in the dark, then irradiated the reaction with 365 nm for ∼30 s to obtain PSS^365^, followed the reaction progress again in the dar**k** for another ∼5 min, then irradiated the reaction with 420 nm for ∼30 s to obtain PSS^420^, and followed the reaction in the dark until it was completed. The activity of ImGPS(hW123AzoF) was increased in PSS^365^ (LRF ∼1.6), remained steady after irradiation, and returned to initial values in PSS^420^ (LRF ∼1.8; **Figure 3A**; **Table S1**) demonstrating reversible control via successive monochromatic irradiation. Likewise, the activity of ImGPS(fS55AzoF) was reduced in PSS^365^ (LRF ∼1.5), remained steady after irradiation and again returned to initial value**s** in PSS^420^ (LRF ∼1.6).

**Figure 3.**
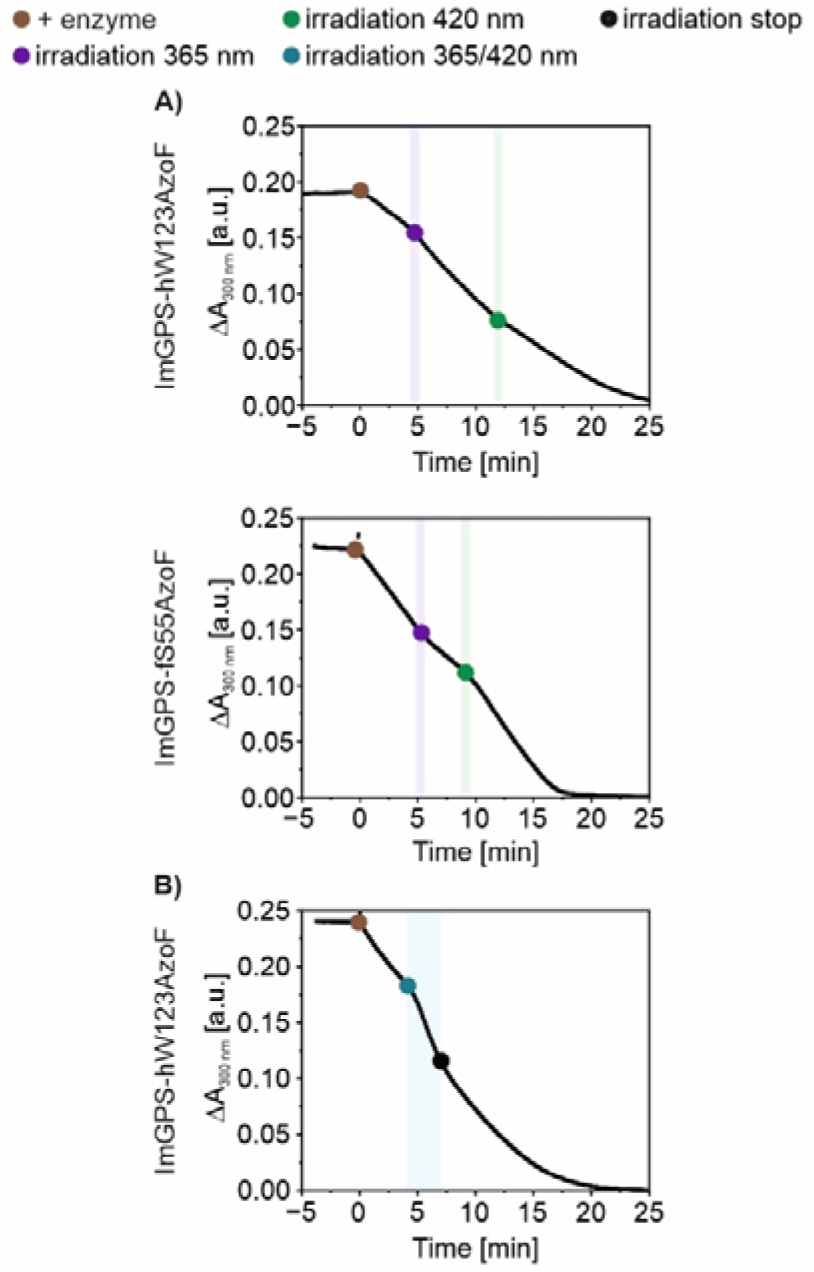
Comparison of ImGPS activity changes facilitated either by successive 365 nm and 420 nm irradiation (A), or by dichromatic 365/420 nm irradiation and subsequent return to the dark (B). For reaction conditions see **Table S1** and **Table S2**.

Next, we measured the effects of dichromatic pulsed irradiation on ImGPS activity with the same experimental se**t**up as described above (**Figure 2C**). Dichromatic irradiation was thereby realized by alternating 365 nm and 420 nm pulses (**Figure S2**) to possibly allow for frequent switching of AzoF between *E*- and *Z*-enriched states. Hence, we started by testing different frequencies of 1–1000 Hz for ImGPS(hW123AzoF), ImGPS(fS55AzoF) as well as wt-ImGPS as a control. The effect of irradiation for each respective variant was the same for all pulsing frequencies. Hence, we summarized the activity and LRF values for all replicates of each variant (“total” in **Table S2**). As a result, we obtained an activity increase ImGPS(hW123AzoF) for with an LRF of ∼1.6. Remarkably, ImGPS(fS55AzoF) and wt-ImGPS remained mainly unaffected by the dichromatic irradiation with LRFs of ∼1.1 and ∼1.0, respectively. We performed further experiments to better assess this result.

First, we were interested whether we could reproduce the behavior of all three variants by simultaneous 365 nm and 420 nm irradiation with LEDs that were powered by a constant current (“constantly powered irradiation”), by which the sample is continuously exposed to light (**Figure S2**). Notably, this experimental setup led to an activity increase not only in ImGPS(hW123AzoF) with an LRF of ∼1.8 but also in ImGPS(fS55AzoF) and wt-ImGPS with LRFs of ∼1.4 and ∼1.2, respectively (**Figure 2D**, grey; **Table S2**) confirming that constantly powered irradiation gives off more heat than pulsed irradiation. To counteract, we repeated our measurements with temperature control set to 20°C. Accordingly, the absolute activity values were smaller in comparison to room temperature (**Figure 2D**, yellow; **Table S2**); however, ImGPS(hW123AzoF) again responded with an activity increase of ∼1.5, whereas ImGPS(fS55AzoF) and wt-ImGPS were nearly unaffected with LRFs of ∼1.1 and ∼1.0, respectively, confirming our initial results. Then, we set out to analyze whether the activity increase during dichromatic irradiation in ImGPS(hW123AzoF) can **b**e primarily associated with the establishment of a new stable *E*:*Z* composition as it was observed for monochromatic irradiati**o**n. For this, we performed measurements with ImGPS(hW123AzoF), in which we limited the pulsed dichromatic irradiation to 3 min and subsequently followed the reaction progress in the dark until it was completed. Remarkably, enzyme activity returned to **t**he initial low activity after the irradiation-facilitated increase with an LRF of ∼2.0 (**Figure 3B**; **Table S2**). Although we cannot exclude a small contribution of the configurational change of AzoF, the observed effect in ImGPS(hW123AzoF) appears to be primarily caused by an alternative mechanism of the photoswitch azobenzene.

We excluded the possibility that the tightly regulated activation of ImGPS(hW123AzoF) during dichromatic irradiation was caused by simple heating of the sample because of three reasons: i) wt-ImGPS showed only insignificant changes in activity during monochromatic and dichromatic irradiation, ii) ImGPS(fS55AzoF) reacted with an activity decrease instea**d** of increase during monochromatic irradiation and was largely inert to dichromatic irradiation, and iii) ImGPS(hW123AzoF) requires a relatively high raise in temperature of Δ*T* ∼5.7–8.4 K to achieve the same activity increase of 1.6–2.0 during dichromatic irradiation (**Figure S3**). Moreover, the consistent LRF generated by different 365 nm and 420 nm pulse frequencies and the fact that temporary activation of ImGPS(hW123AzoF) could be reproduced with simultaneous 365/420 nm irradiation led us to question whether frequent switching of AzoF between *E*-and *Z*-enriched states might be the cause of this effect. To further investigate this possibility, we spectroscopically analyzed the photophysical behavior of AzoF during monochromatic and dichromatic irradiation.

The UV/Vis spectra of hW123AzoF (**Figure S4A**, black) and fS55AzoF (**Figure S5A**, black) in its TEQ^ai^ show a pronounc**e**d peak at ∼280 nm, to which mainly the enzyme environment contributes, and two distinctive peaks at ∼322 nm and ∼425 **n**m representing the _π→π_* and n_→π_* transition of AzoF, respectively. Next, we measured time-resolved UV/Vis spec**t**ra during 365 nm and subsequent 420 nm irradiation (**Figure S4B**,**C**; **Figure S5B**,**C**). By plotting the absorbanc**e** at 322 nm against time we visualized the establishment of **t**he photoinduced *Z*-enriched and *E*-enriched equilibria and determined the overall rates of isomerization *k*^*365*^ and *k*^*420*^ u**s**ing a mono-exponential function as justified else-where.^[12]^ **F**or hW123AzoF (**Figure 4A**), both PSS^365^ and PSS^420^ **we**re established after 20–30 s with *k*^*365*^ ∼ 17.4×10^−2^ s^−1^ and *k*^*420*^ ∼ 21.1×10^−2^ s^−1^, and half-times of *t*_*½*_ ^*365*^ ∼ 4.0 s and *t*_*½*_ ^*420*^ ∼ 3.3 s, respectively. Irradiation of fS55AzoF (**Figure S5D**) was similarly fast with *k*^*365*^ ∼ 21.0×10^−2^ s^−1^ and *k*^*420*^ ∼ 25.6×10^−2^ s^−1^, and half-times of *t*_*½*_ ^*365*^ ∼ 3.3 s and *t*_*½*_ ^*420*^ ∼ 2.7 s, respectively. This confirmed that the PSS^365^ and PSS^420^ were fully establishe**d** by successive monochromatic irradiation in our previous *E*:*Z* ratios and at PSS^420^ of 98*E*:2*Z* for hW123AzoF and 100*E*:0*Z* for during monochromatic irradiation by peak deconvolution with Gaussian functions (**Extended Text S1** and **Figure S6**).^[13^

**Figure 4.**
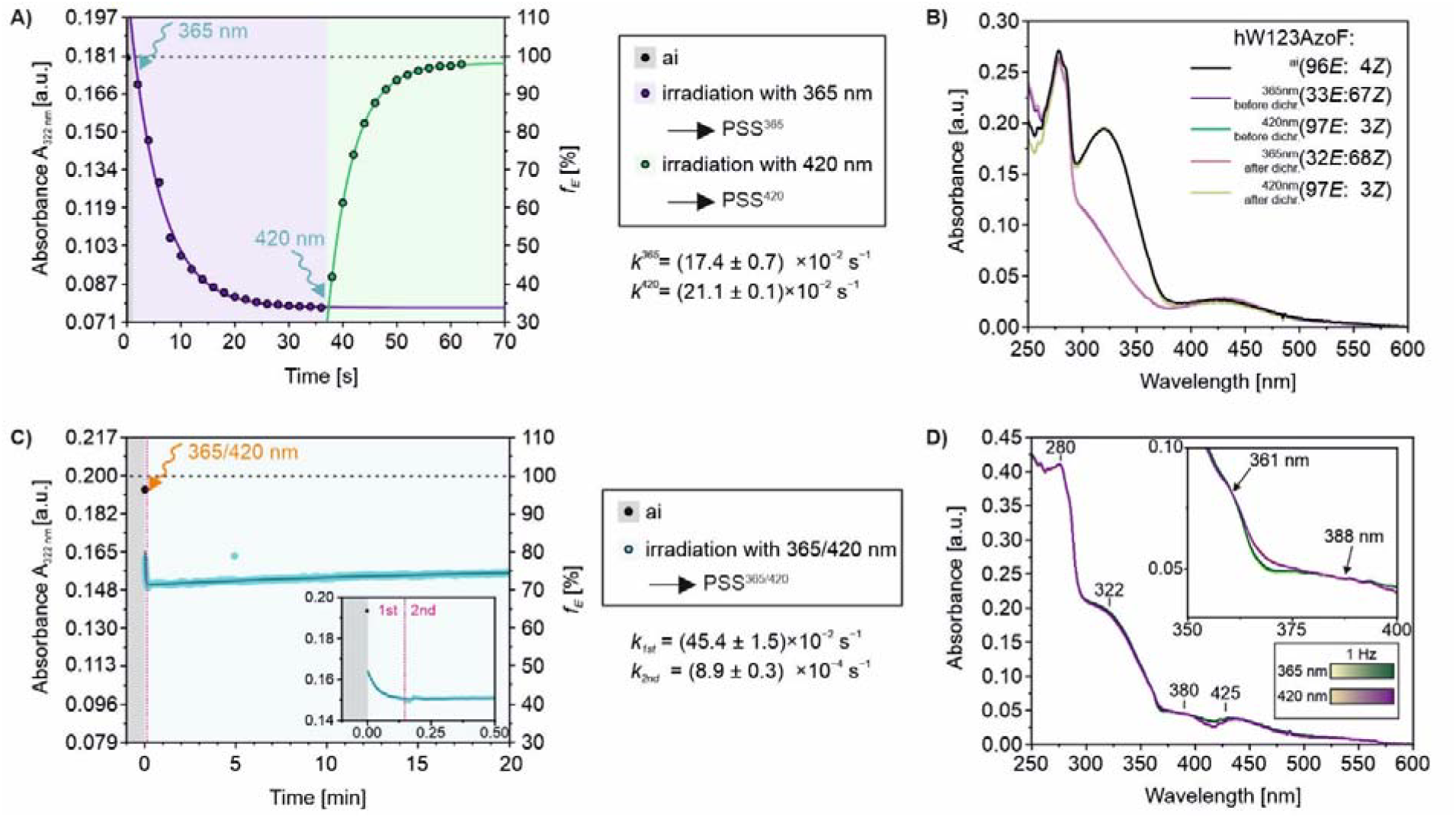
UV/Vis analysis before, after and during irradiation of hW123AzoF. B) Progress curve following the change of the approximate *E* isomer fraction (*f*_*E*_) at 322 nm during 365 nm and subsequent 420 nm constantly powered irradiation. Mono-exponential fitting of each section obtained the rate constants *k*^*365*^ and *k*^*420*^ ± standard error of fit (S.E.). B) Spectra of 15 µM hW123AzoF (in 50 mM HEPES pH 7.5, 100 mM NaCl) in its TEQ^ai^ (black), its PSS^365^, and its PSS^420^. The PSSs were obtained via monochromatic pulsed irradiation either before dichromatic pulsed irradiation (“dichr.”; violet and green) or after (pink and light-green). C) Progress curve following the change of *f*_*E*_ at 322 nm during dichromatic irradiation with 1000 Hz pulse frequency. Double-exponential fitting obtained the rate constants for the first phase *k*_*1st*_ and for the second phase *k*_*2nd*_ ± S.E. D) UV/Vis spectra of 15 µM hW123AzoF acquired with a cycle time of 0.5 s during alternating (1 Hz) 365 nm (green gradient) and 420 nm (violet gradient) pulses. The inset shows an additional isosbestic point of AzoF isomerization at ∼361 nm (arrow) to the one at ∼388 nm (**Figure S4**). See **Extended Text S1** for a description of the approximation of *f*_*E*_.

Since the same LEDs were used, we obtained *E*:*Z* ratios at PSS^365^ of 33*E*:67*Z* for hW123AzoF and 32*E*:68*Z* for fS55AzoF, experiment (**Figure 3A**). We further estimated the^]^ Since the same LEDs were used, we obtained *E*:*Z* ratios at fS55AzoF with both constantly powered (**Figure S4A**; **Figure S5A**) and pulsed (**Figure 4B**; **FigureS7B**) irradiation.

Having determined the photophysical behavior of hW123AzoF and fS55AzoF during monochromatic irradiation, we were interested in the effects of dichromatic irradiation. To this end, we initially measured time-resolved UV/Vis spectra for hW123AzoF during irradiation with various pulse frequencies. Since the rates of isomerization were <1 s^−1^ (**Figure 4A**), we chose 1, 10, 100 and 1000 Hz (**Figure S8A–D**). Intriguingly, the absorbance for 1 Hz irradiation alternated between higher and lower absorbance values with a Δ*A*_*322 nm*_ of ∼0.003 a.u. indicating that a fraction of AzoF switched back and forth in response to the alternating 365 nm and 420 nm irradiation. To estimate the percentage of AzoF that switched with each cycle, we compared the spectra to a cycle performance of hW123AzoF, in which AzoF was switched 31 times from PSS^365^ to PSS^420^ and back resulting in a Δ*A*_*322 nm*_ of ∼0.098 a.u. (**Figure S8E**). Comparison of both Δ*A*_*322 nm*_ showed that the toggling of absorbance in the 1 Hz irradiation experiment was negligible. Hence, AzoF was not able to switch between *E*- and *Z*-enriched states within one second or less confirmed by the 10–1000 Hz irradiation experiments, in which the absorbance signals did not chan**g**e over time. With this, we disproved our initial theory that dichromatic irradiation might lead to frequent switching of AzoF, which then introduces e.g. kinetic energy into the enzyme scaffold stimulating allostery dynamics and enzyme catalysis.

Interestingly, dichromatic irradiation of any frequency resulted in a drop in absorbance at 322 nm of ∼0.033 a.u. compared to the absorbance of TEQ^ai^ (**Figure S8F)** indicating that a photo-induced equilibrium was reached prior to the start of our measurement. To capture the establishment of this equilibrium, we measured time-resolved UV/Vis spectra for hW123AzoF (**Figure S7A**), for which we simultaneously started the measurements and the dichromatic pulses using 1000 Hz and followed the reaction for 20 min. By plotting the absorbance at 322 nm against time we could visualize the formation of a PSS for the dichromatic irradiation. Contrary to monochromatic irradiation, the isomerization reaction the of hW123AzoF did not follow a mono-exponential function (**Figure 4C**). Instead, an initial rapid decline of the signal was followed by a much slower increase. The presence of these two reaction phases indicated that the reaction might follow a double-exponential function, which we used to fit our data. The fit obtained excellent p-values for both rates below the 0.05 significance level (5×10^−168^ and 3×10^−179^), and R^2^ values >0.95. The first reaction occurred with a rate *k*_*1st*_ of ∼45.4×10^−2^ s^−1^ and a half-time of ∼1.5 s on the same timescale as the isomerization reactions induced by monochromatic irradiation. Accordingly, the first phase is concluded after <30 s, which is comparable to the monochromatic irradiation experiments (**Figure 4C** inset and **Figure 4A**). However, the second reaction was >500-fold slower with *k*_*2nd*_ ∼8.9×10^−4^ s^−1^ and a half-time of ∼13.0 min. We also performed the same measurement for fS55AzoF (**Figure S7B– D**), which followed the same general reaction course. However, the first phase was faster than for hW123AzoF, so that we were not able to fit our data with a double-exponential function. Mono-exponential fitting of the second reaction phase obtained an apparent *k*_*2nd*_ of ∼14.8×10^−4^ s^−1^ with a half-time of ∼7.8 min, which is similar to the *k*_*2nd*_ of hW123AzoF. As a result of the double-exponential behavior, hW123AzoF and fS55AzoF exhibited *E*:*Z* ratios of 71*E*:29*Z* and 65*E*:35*Z* after the first phase (PSS1^365/420^), and appeared to approach *E*:*Z* ratios of 75*E*:25*Z* and 73*E*:27*Z* in the second phase (PSS2^365/420^), respectively. Interestingly, PSS2^365/420^ of both enzymes could again be converted to the *Z*-enriched PSSs and *E*-enriched PSSs with monochromatic irradiation (**Figure 4B**; **Figure S7B**).

Intrigued by this unique behavior during dichromatic irradiation, we performed a detailed evaluation to assess whether the double-exponential shape of the progress curve could be associated with the instantly reversible photocontrol in our previous experiments. The composition of the PSS obtained by monochromatic irradiation depends on two isomerization rates, e.g., and (**Figure 1B**). Together they define the overall rate of isomerization, e.g., *k*^*365*^ (**Equation 1**),^[12]^

**Equation 1**

where *d* is the path length of irradiation, *I*_*0*_ is the intensity of the light source and *F* is the photokinetic factor, which depends on the absorbance at the irradiation wavelength. Likewise, the composition of a PSS between *E* and *Z* established by dichromatic irradiation combines four isomerization rates (**Equation 2**).

**Equation 2**

For this equilibrium reaction, we expect a mono-exponential decay of the *E* isomer as observed for PSS1^365/420^. Moreover, the overall rate of isomerization *k*^*365/420*^ should be almost equal to the sum of the two rates *k*^*365*^ and *k*^*420*^ as derived from Equation 1 (**Equation 3**).

**Equation 3**

Looking at the first exponential phase that establishes PSS1^365/420^ of hW123AzoF (**Figure 4C**), we found that its rate constant *k*_*1st*_ is with ∼45.4×10^−2^ s^−1^ very similar to the sum of *k*^*365*^ and *k*^*420*^ of hW123AzoF obtained by monochromatic irradiation (∼38.6×10^−2^ s^−1^; cf. **Figure 4A**). We therefore assigned the initial rapid decline of the progress curve to the isomerization equilibrium between *E* and *Z*.

In the next step, we tried to identify the obvious secondary reaction that took place upon dichromatic irradiation and that caused the double-exponential behavior. For this, we reanalyzed all time-resolved UV/Vis data (**Figure S4–S8**) and found to our astonishment a fourth signal with an absorbance maximu**m** at ∼380 nm that only appeared during dichromatic irradiation alongside the protein signal (∼280 nm), the _π→π_* transition (∼322 nm) and the n_→π_* transition (∼425 nm). This signal was particularly noticeable in the UV/Vis trace of hW123AzoF during 1 Hz irradiation (**Figure 4D**). Because the absorbance alternated between higher and lower absorbance values, we could identify an additional isosbestic point for AzoF isomerization at ∼361 nm during 1 Hz dichromatic irradiation compared to the two at ∼278 nm and ∼388 nm during monochromatic irradiation (**Figure 4D**; **Figure S4**). This confirmed the existence of at least one other species in the reaction. Remarkably, termination of the dichromatic irradiation and subsequent monochromatic irradiation led to the disappearance of this additional UV/Vis signal (**Figure 4B**). This finding indicated that the unknown species is derived from *E* or *Z* AzoF and reacts back to eit**h**er isomer. It further explains the apparent slow increase of the *E* fraction (*f*_*E*_) in **Figure 4C**. As a consequence, the composition of the PSS during dichromatic irradiation will also depend on **t**he reaction rates of this additional secondary reaction and it might include an equilibrium reaction with this apparent third species. We suspect that the signal originates from an isomerization intermediate, however, we are uncertain of the type of transition this intermediate undergoes and why it only occurs during dichromatic irradiation. In a first attempt to identify this intermediate, we performed NMR experiments (**Extended Text S2** and **Figures S9**–**Figure S16**). However, our results indicated that this AzoF species exclusively appears at a defined composition of wavelengths and requires finely tuned light intensities of both LEDs. This represents a challenge for NMR experiments, since the current setups for online irradiation are not equipped for a larger wavelength screen and light intensity is significantly lost in the glass fiber connecting LED and sample tube. Future studies should reveal, which wavelength compositions and which intensities of both LEDs are required to manifest the third AzoF species, which might allow detection via NMR with an improved technical irradiation device.

Obviously, the appearance of the third AzoF species during dichromatic irradiation and its disintegration after return to the dark correlated with the increase and reestablishment of enzyme activity in ImGPS(hW123AzoF). We suspect that the appearance of this intermediate is associated with transient conformational transitions that are essential for enzymatic catalysis and that mediate instantly reversible photocontrol, though we cannot exclude other mechanisms such as vibrational energy transfer. Notably, the AzoF species was detectable in both ImGPS variants, however only ImGPS(hW123AzoF) reacted to dichromatic irradiation with an activity increase. Previous studies have shown that position hW123 is strongly involved as hinge residue in the allosterically important interface closure of the HisH and HisF subunit,^[8]^ and is a central residue within the allosteric network as shown previously by a shortest pathmap analysis.^[14]^ In contrast, position fS55 is localized at the HisF active site and is only close to the allosterically important active site loop 1 in HisF,^[10]^ likely explaining why dichromatic irradiation was not able to photocontrol ImGPS(fS55AzoF).

## Conclusion

Photoswitches have hitherto been employed for the reversible regulation of enzyme activity via monochromatic irradiation. In this study, we have explored the effects of dichromatic irradiation on the prominent photoswitch azobenzene. We have used two previously engineered ImGPS variants that could be effectively regulated by light through incorporation of azobenzene as the unnatural amino acid AzoF. Enzyme kinetic studies and photophysical characterization revealed that dichromatic irradiation followed a different modus operandi than monochromatic irradiation. In particular, the activity of ImGPS(hW123AzoF) was increased during 365/420 nm irradiation and returned to initial values in the dark. In comparison, its activity was also increased with 365 nm irradiation but could only be reestablished by 420 nm irradiation. Moreover, while ImGPS(fS55AzoF) was largely inert to dichromatic irradiation, it could be photocontrolled by monochromatic irradiation. These effects during dichromatic irradiation might be associated with the occurrence of a third transient AzoF species besides the known *E* and *Z* isomer. Importantly, these effects were obtained by alternating 365 nm and 420 nm pulses as well as simultaneous 365/420 nm irradiation, which is more easily implemented.

We considered the potential benefits of particularly the instant return to initial activity values as well as the inertness to photocontrol in certain incorporation positions of AzoF. Dichromatic irradiation could, e.g., ease the sequential photocontrol of different enzymes within a certain application, such as cell studies or biocatalytic multi-enzyme cascades. Using two different photoswitches for this purpose is critical since the wavelengths of isomerization overlap for a great number of photoswitch pairs. However, combination of a photoswitch that is regulated by monochromatic irradiation in one enzyme and a photoswitch that is controlled by dichromatic irradiation in a different enzyme might establish the required specificity of photocontrol. In this regard, further studies will show whether a specific wavelength composition is required for this modus operandi to work, which could further increase **t**he specificity of photocontrol. In this context, analyzing **t**he requirements for the position of AzoF incorporation for this different modus operandi might contribute to the development of a broadly impactful technique.

## Supporting Information

The authors have cited additional references within the Supporting Information.^[15]^

## Acknowledgements

The authors thank Prof. Dr. Remco Sprangers, Prof. Dr. Reinhard Sterner, Dr. Kristina Straub and Prof. Dr. Till Rudack for lively discussions on the topic, help with NMR measurements and proof-reading the manuscript. Most of all, we are grateful for the strong technical support team consisting of Julia Zach, Jeannette Ueckert and Achim Stark. Finally, we like to thank all our reviewers for their valuable expertise and helpful critique.

## Entry for the Table of Contents

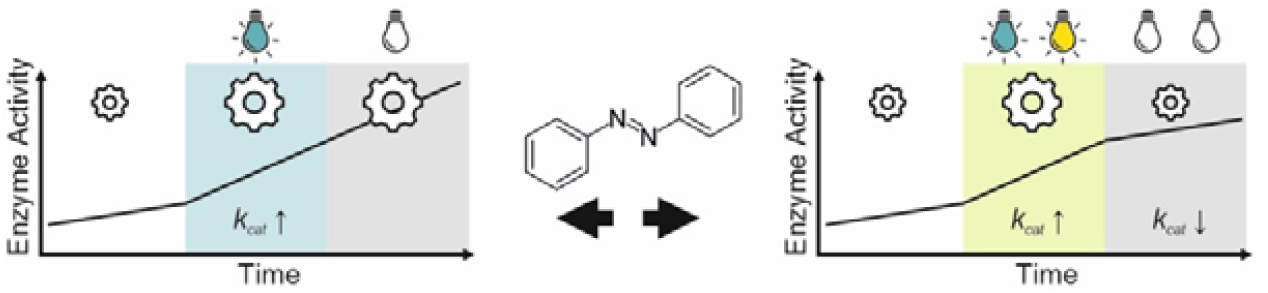

Light induced activation of enzyme activity with azobenzene incorporated as unnatural amino acid is stable after monochromatic irradiation and instantly reversible upon dichromatic irradiation highlighting a hitherto undiscovered modus operandi.

## References

[1] a)A. C. Kneuttinger, Biol. Chem. 2022, 403, 573; b) M. M. Lerch, M. J. Hansen, G. M. van Dam, W. Szymanski, B. L. Feringa, Angew. Chem., Int. Ed. 2016, 55, 10978; c) K. Hüll, J. Morstein, D. Trauner, Chem. Rev. 2018, 118, 10710; d) W. Szymański, J. M. Beierle, H. A. V. Kistemaker, W. A. Velema, B. L. Feringa, Chem. Rev. 2013, 113, 6114.

[2] a)C. J. Brown, Acta Crystallogr. 1966, 21, 146; b) C. R. Crecca, A. E. Roitberg, J. Phys. Chem. A 2006, 110, 8188.

[3] C. L. Forber, E. C. Kelusky, N. J. Bunce, M. C. Zerner, J. Am. Chem. Soc. 1985, 107, 5884.

[4] a)L. Gagliardi, G. Orlandi, F. Bernardi, A. Cembran, M. Garavelli, Theor. Chem. Acc. 2004, 111, 363; b) A. Mostad, C. Rømming, S. Hammarström, R. J. J. C. Lousberg, U. Weiss, Acta Chem. Scand. 1971, 25, 3561.

[5] V. Ladányi, P. Dvořák, J. Al Anshori, L. Vetráková, J. Wirz, D. Heger, Photochem. Photobiol. Sci. 2017, 16, 1757.

[6] M. Bose, D. Groff, J. Xie, E. Brustad, P. G. Schultz, J. Am. Chem. Soc. 2006, 128, 388.

[7] A. C. Kneuttinger, K. Straub, P. Bittner, N. A. Simeth, A. Bruckmann, F. Busch, C. Rajendran, E. Hupfeld, V. H. Wysocki, D. Horinek et al., Cell Chem. Biol. 2019, 26, 1501–1514.

[8] A. C. Kneuttinger, C. Rajendran, N. A. Simeth, A. Bruckmann, B. König, R. Sterner, Biochemistry 2020, 59, 2729.

[9] A. Douangamath, M. Walker, S. Beismann-Driemeyer, M. Vega-Fernandez, R. Sterner, M. Wilmanns, Structure 2002, 10, 185.

[10] S. Beismann-Driemeyer, R. Sterner, J. Biol. Chem. 2001, 276, 20387.

[11] C. Hiefinger, S. Mandl, M. Wieland, A. Kneuttinger in Methods in Enzymology (Ed.: A. K. Shukla), Academic Press, 2023, pp. 247–288.

[12] E. Stadler, A. Eibel, D. Fast, H. Freißmuth, C. Holly, M. Wiech, N. Moszner, G. Gescheidt, Photochem. Photobiol. Sci. 2018, 17, 660.

[13] a)K. Rustler, P. Nitschke, S. Zahnbrecher, J. Zach, S. Crespi, B. König, J. Org. Chem. 2020, 85, 4079; b) N. A. Simeth, T. Kinateder, C. Rajendran, J. Nazet, R. Merkl, R. Sterner, B. König, A. C. Kneuttinger, Chem. Eur. J. 2021, 27, 2439; c) J. Calbo, C. E. Weston, A. J. P. White, H. S. Rzepa, J. Contreras-García, M. J. Fuchter, J. Am. Chem. Soc. 2017, 139, 1261.

[14] C. Calvó-Tusell, M. A. Maria-Solano, S. Osuna, F. Feixas, J. Am. Chem. Soc. 2022, 144, 7146.

[15] a)W. L. Butler, S. B. Hendricks, H. W. Siegelman, Photochem. Photobiol. 1964, 3, 521; b) J. Luo, S. Samanta, M. Convertino, N. V. Dokholyan, A. Deiters, ChemBioChem 2018, 19, 2178; c)C. Feldmeier, H. Bartling, E. Riedle, R. M. Gschwind, J. Magn. Reson. 2013, 232, 39; d)V. J. Davisson, I. L. Deras, S. E. Hamilton, L. L. Moore, J. Org. Chem. 1994, 59, 137; e) T. J. Klem, V. J. Davisson, Biochemistry 1993, 32,5177; f)F. List, M. C. Vega, A. Razeto, M. C. Häger, R. Sterner, M. Wilmanns, Chem. Biol. 2012, 19, 1589; g) J. P. Wurm, S. Sung, A. C. Kneuttinger, E. Hupfeld, R. Sterner, M. Wilmanns, R. Sprangers, Nat. Commun. 2021, 12, 2748; h) J. M. Lipchock, J. Patrick Loria, Biomol. NMR Assign. 2008, 2, 219; i) J. Lipchock, J. P. Loria, J. Biomol. NMR 2009, 45, 73.

