## Supporting Information for "Instantly Reversible Enzyme Regulation via Dichromatic Irradiation of Incorporated Azobenzene"

[c] F. Zequiri
Department of Biophysics
Ruhr University Bochum
Universitätsstraße 150, D-44780 Bochum (Germany)

†The authors contributed equally

### Table of Contents

Supporting Material ... 3
Figure S1. ... 3
Figure S2. ... 3
Table S1. ... 4
Table S2. ... 5
Figure S3. ... 6
Figure S4. ... 6
Figure S5. ... 7
Extended Text S1 ... 7
Figure S6. ... 8
Figure S7. ... 9
Figure S8. ... 10
Extended Text S2 ... 11
Figure S9. ... 12
Figure S10. ... 12
Figure S11. ... 13
Figure S12. ... 13
Figure S13. ... 14
Figure S14. ... 14
Figure S15. ... 15
Figure S16. ... 15
Experimental Section ... 16
Chemicals and other resources ... 16
Table S3. Key resources ... 16
Irradiation setups ... 16
Table S4. Irradiation setups used in this work. ... 17
Synthesis of AzoF ... 17
Synthesis of ProFAR ... 18
Auxiliary enzymes ... 18
Expression and purification of HisF, HisH, hW123AzoF, fS55AzoF, and $^{15}$N-fF23AzoF ... 18
UV/Vis analysis ... 19
Estimation of the *E*:*Z* ratio from UV/Vis spectra ... 19
Determination of ImGPS activity in the dark and during irradiation ... 20
Nuclear magnetic resonance (NMR) spectroscopy of AzoF ... 20
References ... 21

#### SUPPORTING MATERIAL

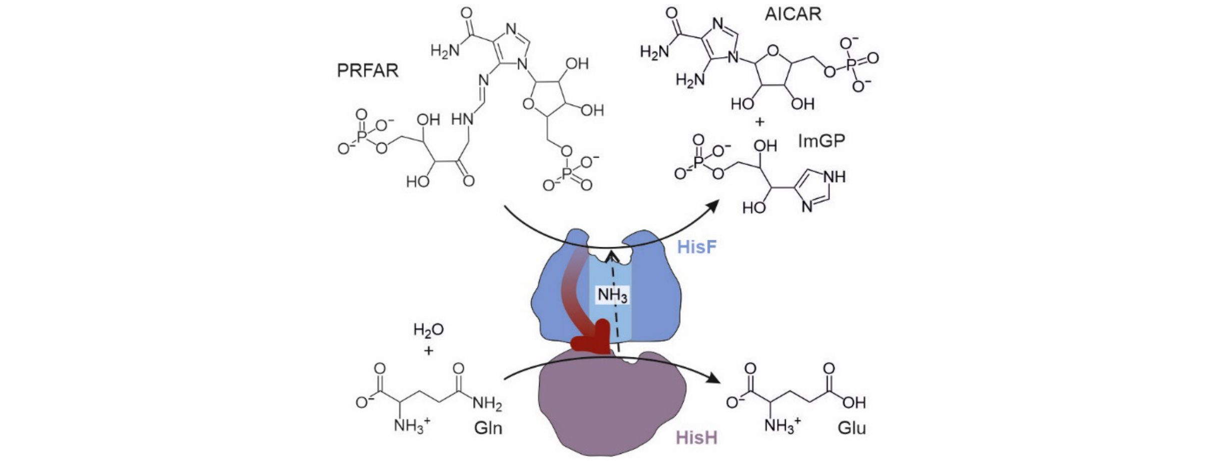


**Figure S1.** Reaction catalyzed by HisH and HisF in the ImGPS complex. HisH (violet) is allosterically activated (red arrow) by binding of PRFAR to the HisF active site and catalyzes the hydrolysis of glutamine (Gln) to glutamate (Glu) and ammonia. The latter travels through an ~25 Å long intermolecular channel (light blue) to the active site of HisF (blue) to react with the HisF substrate PRFAR yielding AICAR and ImGP. PRFAR: N'-[(5'-phosphoribulosyl)formimino]-5-aminoimidazole-4-carboxamide ribonucleotide; AICAR: 5-aminoimidazol-4-carboxamidribotide; ImGP: imidazole glycerol phosphate; Gln: glutamine; Glu: glutamate.

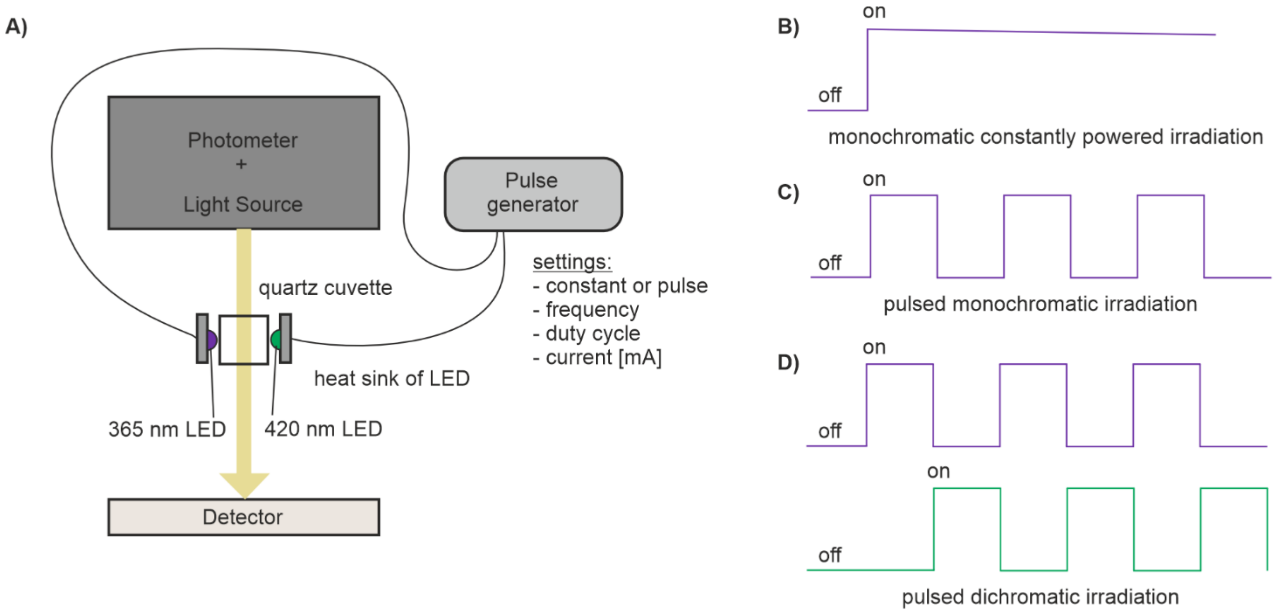


**Figure S2.** Schematic illustration of the UV/Vis irradiation setup and the possible irradiation modes. A) The UV/Vis setup allowed for the monochromatic or dichromatic irradiation with 365 nm and 420 nm using either a constant current ("constantly powered irradiation") or pulse width modulation ("pulsed irradiation"). The LEDs (cf. Table S1) were installed perpendicular to the UV/Vis spectrophotometer measurement beam to enable irradiation of the sample while measuring simultaneously. Adjustable parameters include: current (100-800 mA), frequency (1-10,000 Hz), and duty cycle (10-100%). B–D) Schematic voltage profile of monochromatic constantly powered irradiation (B), monochromatic pulsed irradiation (C), and dichromatic pulsed irradiation (D).

**Table S1. ImGPS activities subjected to monochromatic irradiation.**

| Wavelength | Variant | $k_{cat,app}$*(dark)* [min$^{-1}$] | $k_{cat,app}$*(during hv)* [min$^{-1}$] | $k_{cat,app}$*(after hv)* [min$^{-1}$] | LRF[a] | Replicates |
|---|---|---|---|---|---|---|
| **During irradiation** | | | | | | |
| 365 nm | ImGPS(hW123AzoF) | 23.0 ± 2.2 | 42.4 ± 5.9 | — | 1.86 ± 0.23 ↑ | 4 |
| | ImGPS(fS55AzoF) | 11.8 ± 2.3 | 7.0 ± 1.3 | — | 1.68 ± 0.02 ↓ | 3 |
| | wt-ImGPS | 53.6 ± 5.9 | 52.8 ± 5.2 | — | 1.01 ± 0.02 ↓ | 3 |
| 420 nm | ImGPS(hW123AzoF) | 24.1 ± 3.4 | 23.8 ± 3.1 | — | 1.01 ± 0.03 ↓ | 3 |
| | ImGPS(fS55AzoF) | 12.3 ± 1.4 | 14.3 ± 1.5 | — | 1.17 ± 0.07 ↑ | 6 |
| | wt-ImGPS | 53.3 ± 1.9 | 53.0 ± 3.5 | — | 1.01 ± 0.04 ↓ | 3 |
| **After 30 s irradiation** | | | | | | |
| 365 nm | ImGPS(hW123AzoF) | 23.3 ± 2.2 | — | 37.2 ± 4.6 | 1.60 ± 0.08 ↑[b] | 4 |
| 420 nm | | | — | 20.3 ± 1.5 | 1.83 ± 0.18 ↓[c] | |
| 365 nm | ImGPS(fS55AzoF) | 13.9 ± 1.3 | — | 9.5 ± 1.3 | 1.47 ± 0.06 ↓[b] | 4 |
| 420 nm | | | — | 15.6 ± 2.3 | 1.63 ± 0.12 ↑[c] | |

ImGPS activity: 50 mM Tris-acetate (pH 7.5), 0.6 μM HisA (to convert ProFAR to PRFAR), ~40 μM (saturated) ProFAR, 10 mM (saturated) glutamine, and 0.025–0.2 mM ImGPS at rt. [a] The activity change is indicated by an upwards (activity increase) or downwards arrow (activity decrease). [b] LRF compares $k_{cat,app}$*(dark)* and $k_{cat,app}$*(365 nm)*. [c] LRF compares $k_{cat,app}$*(365 nm)* and $k_{cat,app}$*(420 nm)*. Note: variations of the activity in the dark can be attributed to the use of biological samples. Note: $k$ and LRF values are given as mean ± S.E.M. summarizing the individual values of all replicates.

**Table S2. ImGPS activities subjected to dichromatic irradiation.**

| Variant | Pulse frequency | $k_{cat,app}$(*dark*) [$min^{-1}$] | $k_{cat,app}$(*during hv*) [$min^{-1}$] | $k_{cat,app}$(*after hv*) [$min^{-1}$] | LRF[a] | Replicates |
|---|---|---|---|---|---|---|
| **Pulsed irradiation at room temperature[b]** | | | | | | |
| ImGPS(hW123AzoF) | 1 Hz | 20.6 ± 3.0 | 31.7 ± 5.0 | — | 1.54 ± 0.15 ↑ | 4 |
| | 10 Hz | 19.6 ± 1.8 | 30.7 ± 3.0 | — | 1.57 ± 0.14 ↑ | 4 |
| | 100 Hz | 22.9 ± 2.8 | 37.8 ± 6.0 | — | 1.63 ± 0.24 ↑ | 5 |
| | 1000 Hz | 24.4 ± 2.0 | 38.2 ± 3.5 | — | 1.57 ± 0.16 ↑ | 10 |
| | **total** | **21.1 ± 1.4** | **33.6 ± 2.6** | **—** | **1.59 ± 0.05 ↑** | **23** |
| ImGPS(fS55AzoF) | 1 Hz | 7.8 ± 1.0 | 7.9 ± 1.3 | — | 1.02 ± 0.10 ↓ | 5 |
| | 10 Hz | 10.1 ± 0.9 | 9.2 ± 1.2 | — | 1.11 ± 0.11 ↓ | 4 |
| | 100 Hz | 8.6 ± 1.3 | 8.1 ± 1.2 | — | 1.07 ± 0.05 ↓ | 6 |
| | 1000 Hz | 11.3 ± 1.2 | 11.3 ± 1.6 | — | 1.04 ± 0.08 ↓ | 7 |
| | **total** | **8.7 ± 0.7** | **8.4 ± 0.7** | **—** | **1.05 ± 0.03 ↓** | **22** |
| wt-ImGPS | 1 Hz | 53.5 ± 5.1 | 55.3 ± 7.2 | — | 1.03 ± 0.05 ↑ | 3 |
| | 10 Hz | 55.2 ± 8.4 | 52.3 ± 1.8 | — | 1.05 ± 0.12 ↓ | 2 |
| | 100 Hz | 65.5 ± 3.3 | 63.4 ± 10.6 | — | 1.05 ± 0.12 ↓ | 2 |
| | 1000 Hz | 63.5 ± 5.7 | 61.2 ± 6.0 | — | 1.05 ± 0.05 ↓ | 7 |
| | **total** | **60.4 ± 3.3** | **59.0 ± 3.5** | **—** | **1.03 ± 0.03 ↓** | **14** |
| **Constantly powered irradiation at room temperature[b]** | | | | | | |
| ImGPS(hW123AzoF) | — | 18.1 ± 1.9 | 32.6 ± 3.7 | — | 1.80 ± 0.11 ↑ | 5 |
| ImGPS(fS55AzoF) | — | 10.0 ± 4.2 | 13.9 ± 6.4 | — | 1.38 ± 0.23 ↑ | 3 |
| wt-ImGPS | — | 50.9 ± 5.6 | 60.4 ± 4.6 | — | 1.20 ± 0.07 ↑ | 3 |
| **Constantly powered irradiation at 20°C[c]** | | | | | | |
| ImGPS(hW123AzoF) | — | 13.4 ± 0.6 | 20.3 ± 3.5 | — | 1.50 ± 0.23 ↑ | 5 |
| ImGPS(fS55AzoF) | — | 3.9 ± 0.3 | 3.6 ± 0.4 | — | 1.11 ± 0.08 ↓ | 5 |
| wt-ImGPS | — | 22.9 ± 2.3 | 23.1 ± 1.7 | — | 1.02 ± 0.03 ↑ | 5 |
| **3 min pulsed irradiation and recovery in the dark** | | | | | | |
| ImGPS(hW123AzoF) | 1000 Hz | 23.4 ± 0.8 | 46.8 ± 3.1 | | 2.00 ± 0.26 ↑ | 6 |
| | | | 46.8 ± 3.1 | 23.5 ± 1.6 | 2.04 ± 0.48 ↓ | |

ImGPS activity: 50 mM Tris-acetate (pH 7.5), 0.6 µM HisA (to convert ProFAR to PRFAR), ~40 µM (saturated) ProFAR, 10 mM (saturated) glutamine, and 0.025–0.2 mM ImGPS. [a] The activity change is indicated by an upwards (activity increase) or downwards arrow (activity decrease). [b] Not subjected to temperature control. [c] Subjected to temperature control. Note: variations of the activity in the dark can be attributed to the use of biological samples. Note: *k* and LRF values are given as mean ± S.E.M. summarizing the individual values of all replicates.

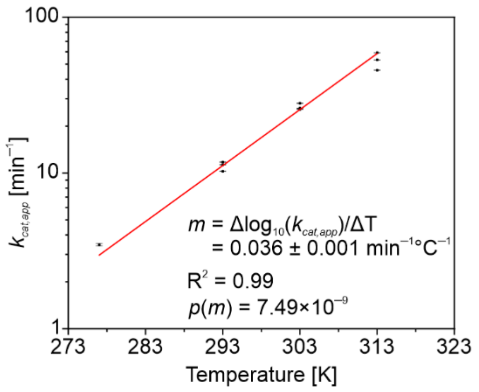


**Figure S3.** Temperature dependence of $k_{cat,app}$ for ImGPS(hW123AzoF) in its as-isolated state. The semi-logarithmic scale allowed for a linear fit obtaining the slope m as the relationship between activity and temperature. A LRF of 1.6–2.0 corresponds to an activity increase of ΔT = m × Δlog(1.6–2.0) = 5.7–8.4 K. ImGPS activity: 50 mM Tris-acetate (pH 7.5), 0.6 µM HisA (to convert ProFAR to PRFAR), ~40 µM (saturated) ProFAR, 10 mM (saturated) glutamine, and 0.1 mM ImGPS.

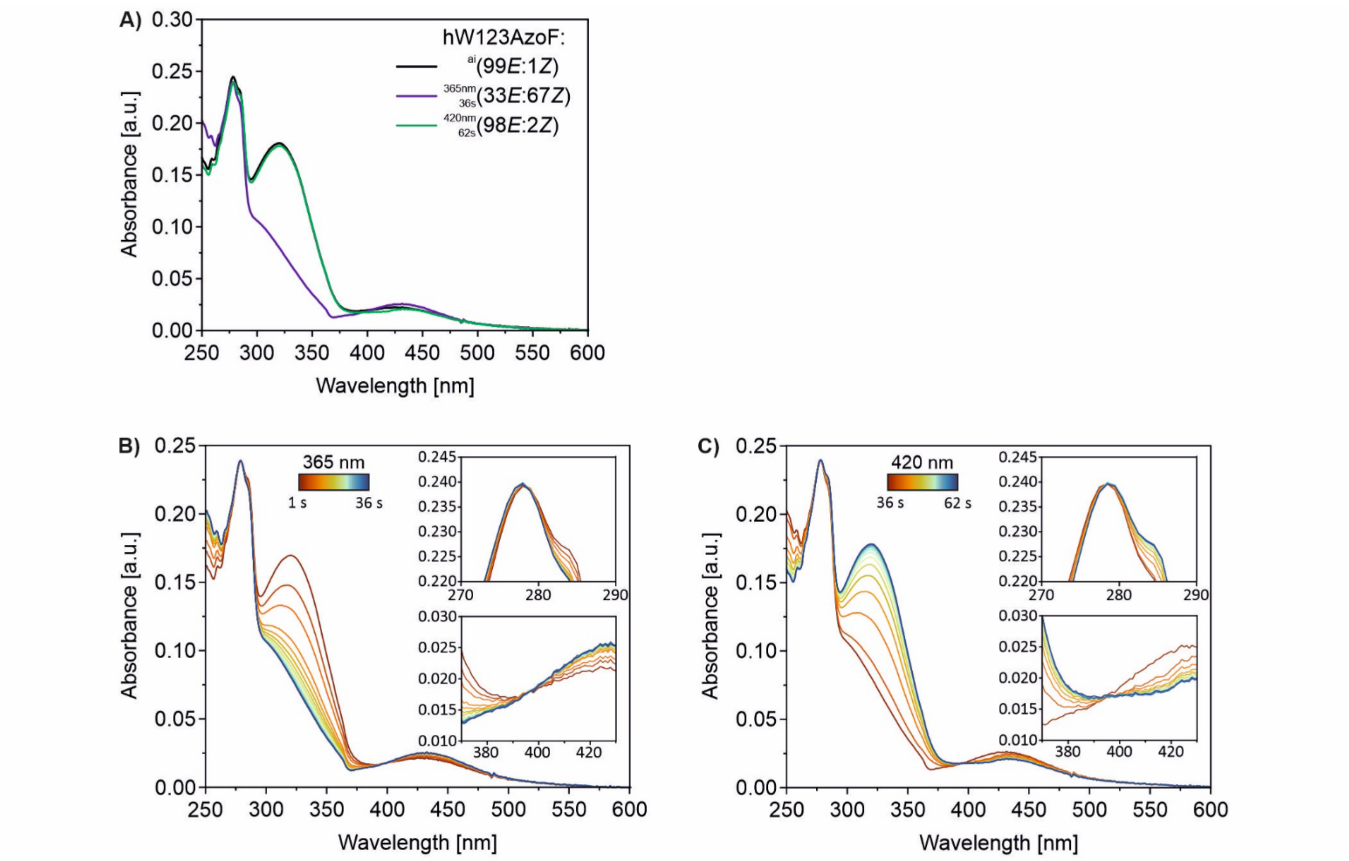


**Figure S4.** Time-resolved UV/Vis analysis of 15 µM hW123AzoF (in 50 mM HEPES pH 7.5, 100 mM NaCl) using monochromatic constantly powered irradiation. A) Spectra of 15 µM hW123AzoF (in 50 mM HEPES pH 7.5, 100 mM NaCl) in its TEQ$^{ai}$ (black), its PSS$^{365}$ (violet), and its PSS$^{420}$ (green) (cf. **Figure 4A**). B) UV/Vis spectra acquired with a cycle time of 2 s during 365 nm irradiation (violet section in **Figure 4A**). C) UV/Vis spectra acquired with a cycle time of 2 s during 420 nm irradiation (green section in **Figure 4A**). The insets in subpanels B and C show the isosbestic points of AzoF isomerization at ~278 nm and ~395 nm.

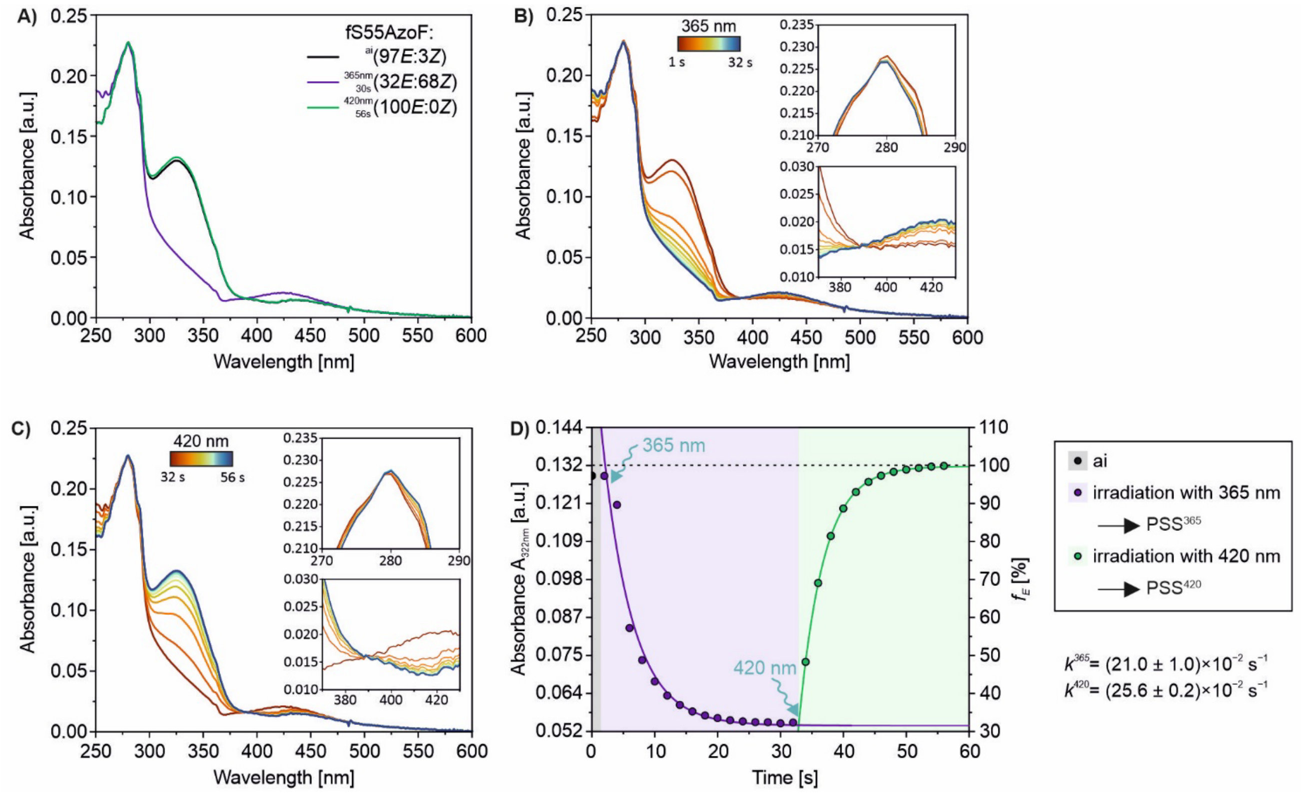


**Figure S5.** UV/Vis analysis before, after and during monochromatic constantly powered irradiation of fS55AzoF. A) Spectra of 15 µM fS55AzoF (in 50 mM HEPES pH 7.5, 100 mM NaCl) ) in its TEQ$^{ai}$ (black), its PSS$^{365}$ (violet), and its PSS$^{420}$ (green) (cf. D). B) UV/Vis spectra acquired with a cycle time of 2 s during 365 nm irradiation (violet section in D). C) UV/Vis spectra acquired with a cycle time of 2 s during 420 nm irradiation (green section in D). The insets in subpanels B and C show the isosbestic points of AzoF isomerization at ~278 nm and ~388 nm. D) Progress curve following the change of the *E* isomer fraction ($f_E$) at ~322 nm during 365 nm and subsequent 420 nm irradiation. See **Figure S6** for a description of the approximation of $f_E$. Mono-exponential fitting of each section obtained the rate constants $k^{365}$ and $k^{420}$ ± S.E..

#### Extended Text S1

For each UV/Vis spectrum, the π→π* and n→π* transitions of AzoF were fitted with the multiple peak fitting tool in Origin 2024 using a Gaussian function, which yielded a cumulative fit ("*Gauss Fit*" in **Figure S5A–C**; dashed black and blue lines). To estimate the *E*:*Z* ratio the 0*E*:100*Z* spectrum (dashed violet line) was simulated using **Equation S5** in which $A_i$ corresponds to the spectra after 4 s, 10 s and 24 s irradiation (blue Gauss Fits), $A_E$ to the spectrum of the as-isolated state (black Gauss Fits) and $f_E$ to the fraction of the *E* isomer. $f_E$ was obtained by manual adjustment until the signal of the π→π* absorbance band approximates zero. $f_E$ values are given in each subpanel of **Figure S5A–C**. $f_E$ values were then determined for multiple spectra in one time-resolved experiment and plotted against the measured absorbance at 322 nm (**Figure S5D**). Finally, the resulting linear equation of the fit was used to determine the approximate fraction of *E* for the as-isolated state and each timepoint during irradiation. Note that our estimation relies on the assumption that the as-isolated protein is thermally equilibrated to 100*E*:0*Z*. However, minimal exposure to light during the handling of the samples can easily disrupt the thermal equilibrium and change the *E*:*Z* ratio introducing a bias into the approach. More quantitative ways of estimation such as high-performance liquid chromatography as used for isolated AzoF,[10] or spectrophotometric determination techniques as used for photoreceptors[11] are not viable for AzoF-containing proteins; for the latter this is due to the completely overlapping absorbance spectra of *E* and *Z*. Hence, slight deviations in *E*:*Z* ratios at the PSS, when compared to other studies,[1,2,10] might be traced back to this bias.

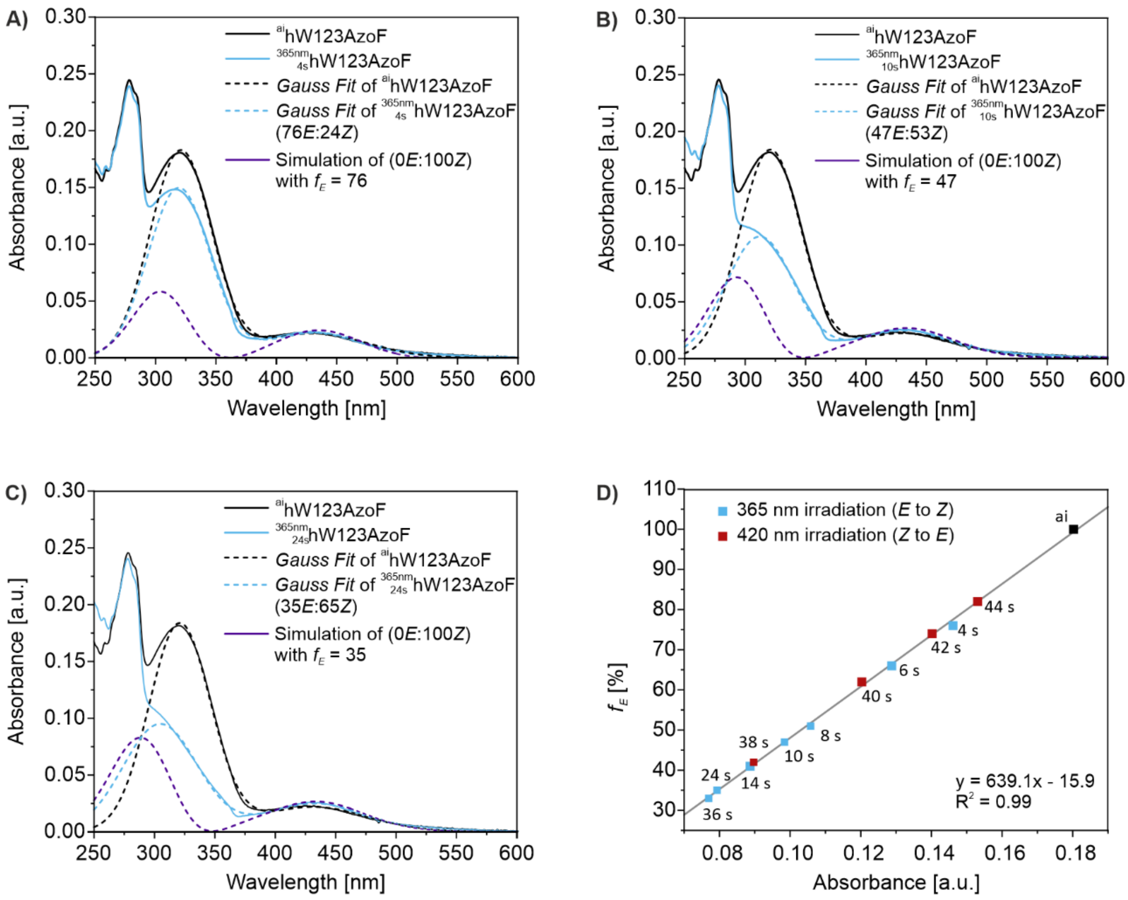


**Figure S6.** Determination of the of *E*:*Z* ratio using peak deconvolution with Gaussian functions shown exemplarily for spectra from **Figure S4**. A–C) UV/Vis spectra of 15 µM hW123AzoF (in 50 mM HEPES pH 7.5, 100 mM NaCl) in its as-isolated state (ai; solid black line) and after 4 s (A), 10 s (B), and 24 s (C) of 365 nm irradiation (solid blue line). Gauss fits and simulations are provided as dashed lines D) Linear fit to determine the fraction of the *E* isomer ($f_E$) from absorbance values at 322 nm.

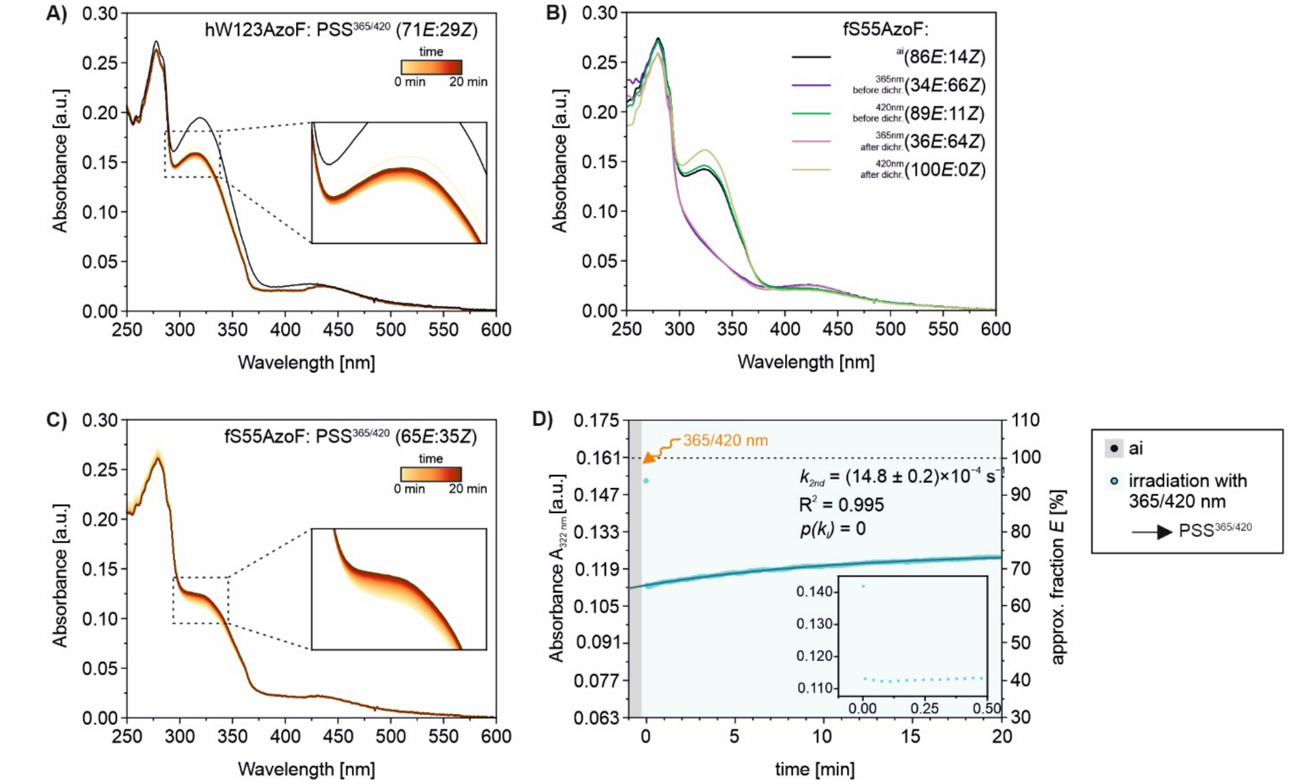


**Figure S7.** Time-resolved UV/Vis analysis of 15 µM hW123AzoF and fS55AzoF (in 50 mM HEPES pH 7.5, 100 mM NaCl) using dichromatic pulsed irradiation at 1000 Hz. A) UV/Vis spectra of hW123AzoF acquired with a cycle time of 0.5 s during irradiation. Note that spectra are shown only for every 2 s. B) UV/Vis spectra of fS55AzoF in its TEQ$^{ai}$(black), PSS$^{365}$ and PSS$^{420}$. The PSSs were obtained via monochromatic pulsed irradiation either before dichromatic pulsed irradiation ("dichr."; violet and green) or after dichromatic pulsed irradiation (pink and light-green). C) UV/Vis spectra of fS55AzoF acquired with a cycle time of 2 s during irradiation. D) Progress curve following the change of the *E* isomer fraction at 322 nm during irradiation. See **Figure S5** for a description of the approximation of $f_E$. A double-exponential fit was not feasible, but mono-exponential fitting obtained the rate constant for the second phase $k_{2nd}$ ± S.E..

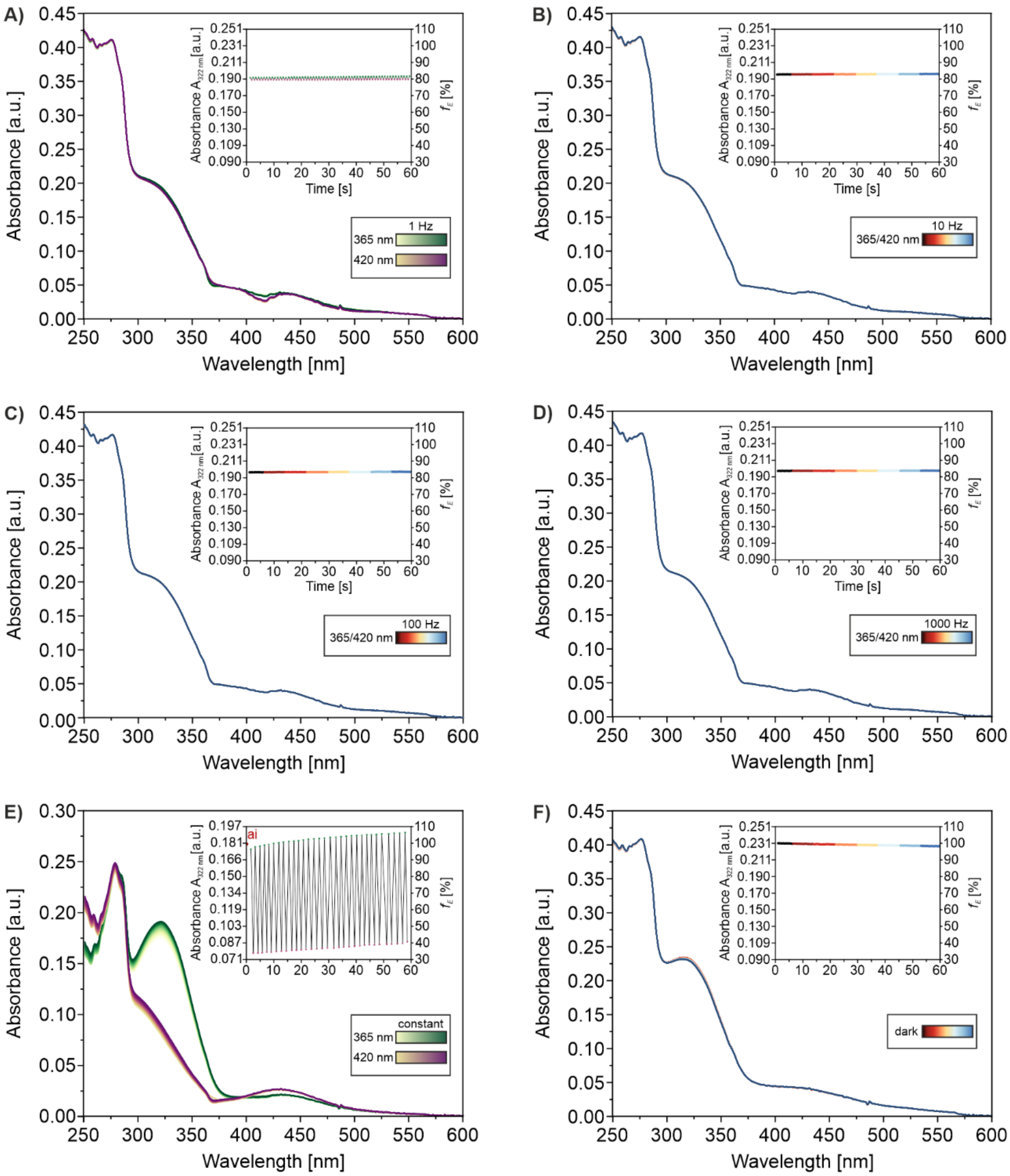


**Figure S8.** Time-resolved UV/Vis analysis of 15 μM hW123AzoF (in 50 mM HEPES pH 7.5, 100 mM NaCl) using dichromatic pulsed irradiation at different pulsing frequencies. A) UV/Vis spectra acquired with a cycle time of 0.5 s during dichromatic irradiation with 1 Hz pulse frequency. Since we used alternating monochromatic pulses, every second spectrum corresponds to 365 nm or 420 nm irradiation as visualized with a green and violet gradient. Inset: Absorbance at 322 nm alternated between higher and lower absorbance values indicating that a fraction of AzoF switched back and forth. B–D) UV/Vis spectra acquired with a cycle time of 0.5 s during dichromatic irradiation with 10 Hz (B), 100 Hz (C) and 1000 Hz (D) pulse frequency. Inset: The cycle time of 0.5 s could not resolve the response to the rapidly changing 365 nm and 420 nm irradiation and the absorbance signal was consistent throughout the measurement indicating that isomerization of AzoF established a photo-induced equilibrium. E) Cycle performance of hW123AzoF. UV/Vis spectra were recorded for 31 cycles of 365 nm and subsequent 420 nm constantly powered irradiation. Only the spectra at the $PSS^{365}$ or $PSS^{420}$ are shown. Inset: Absorbance at 322 nm alternated between higher and lower absorbance values indicating that AzoF switched back and forth between $PSS^{365}$ or $PSS^{420}$. F) Control spectra of hW123AzoF acquired with a cycle time of 0.5 s in the dark. Note: Spectra were recorded only after the start of irradiation.

#### Extended Text S2

We aimed to investigate the third azobenzene species via NMR. To this end, we used a previously established in situ irradiation device for NMR tubes.[4] This setup consists of a power supply, a potentiometer to adjust the current through the LED and a transistor controlled by the spectrometer. The LED output is coupled to an optical fiber with a roughened tip that is inserted in a coaxial inlet inside the NMR tube. Notably, transmittance through the optical fiber reduces the output power of light ~10-fold in the original setup substantiating the need for high power LEDs.[4]

We initially assigned the signals for the *E* and *Z* isomer in 10 mM DMSO-$d_6$ at 25°C using different NMR experiments [1D-$^1$H; 2D-$^1$H, $^1$H correlated spectroscopy (COSY); 2D-$^1$H, $^{13}$C heteronuclear multiple bond correlation (HMBC); 2D-$^1$H, $^{13}$C heteronuclear single quantum coherence (HSQC)] (**Figures S9–S13**). For this, we irradiated 10 mM AzoF with UV light at 25°C until a PSS was reached. We then conducted the NMR experiments of this *E*:*Z* mixture in the dark.

For the excitation of the π→π* transition of isolated AzoF we chose a pre-installed 365 nm high-power LED similar to our UV/Vis analyses. However, we needed to test various LEDs for the excitation of the n→π* transition, since a 420 nm high-power LED was not available. The pre-installed 530 nm high-power LED achieved the best results regarding the isomerization rate and PSS. Both LEDs were thereby powered with a high, constant current of 1.0 A. After ~2 min of 365 nm irradiation a PSS of 16*E*:84*Z* was obtained, whereas 4 min of 530 nm irradiation was required to achieve the second PSS of 69*E*:31*Z*. The respective isomerization rates (**Figure S14**; $k^{365}$ ~$4.9\times10^{-2}$ s$^{-1}$ and $k^{530}$ ~$1.5\times10^{-2}$ s$^{-1}$) were 4–13-fold slower than the rates for hW123AzoF and fS55AzoF in our UV/Vis analyses ($k^{36m}$ ~ $17.4\times10^{-2}$ s$^{-1}$ and ~ $21.0\times10^{-2}$ s$^{-1}$, $k^{420}$ ~ $21.2\times10^{-2}$ s$^{-1}$ and $25.6\times10^{-2}$ s$^{-1}$, respectively).

Since the PSSs were still either *E*- or *Z*-enriched and the isomerization rates were still in a similar range as the ones determined in our UV/Vis analyses, we set out to test both LEDs for NMR studies during dichromatic irradiation. For this purpose, we designed a beta version of the original NMR setup in collaboration with HK Testsysteme GMbH, which allowed the pulsed irradiation with two different wavelengths. Here, we integrated a bifurcated glass fiber that combines two different LED outputs to the single optical fiber inside the NMR tube (**Figure S15**). Moreover, we equipped this setup with a two-channel pulse generator. This way, we could apply monochromatic constantly powered or dichromatic pulsed irradiation as used for our UV/Vis and activity measurements. Since the same LEDs as used in the original setup were no longer available, we chose to use high-power LEDs that were as similar as possible to the ones in the original setup with 365 nm and 524 nm (**Table S2**). We regarded it necessary to choose high-power LEDs, because by inserting the bifurcated glass fiber we might encounter an even greater loss of light intensity as observed in the original setup.

Next, we set out to test this beta version of the irradiation setup with an ImGPS sample containing AzoF. We selected position fF23 for the incorporation of AzoF due to the following three reasons. First, our previous studies have shown that heterologous gene expression and subsequent purification using affinity chromatography obtains relatively high amounts of fF23AzoF (~34.7 mg per liter expression medium),[1] which is a prerequisite for the expression in isotope enriched minimal medium for NMR experiments. Second, NMR spectra of HisF have already been solved and annotated.[12] Third, the NMR signal of position fF23 is well-defined and separated from other signals. Thus, we produced fF23AzoF using heterologous gene expression in minimal medium containing $^{15}NH_4Cl$ for NMR-labeling. To check whether AzoF was properly incorporated we recorded UV/Vis spectra of $^{15}$N-fF23AzoF in its TEQ$^{ai}$, PSS$^{365}$ and PSS$^{420}$ using monochromatic constantly powered irradiation (**Figure S16A**). The *E*:*Z* ratios at PSS$^{365}$ and PSS$^{420}$ of 21*E*:79*Z* as well as 81*E*:19*Z*, respectively, confirmed the successful incorporation and integrity of AzoF, despite a small variation from the PSSs obtained for hW123AzoF and fS55AzoF (cf. **Figure 2A**; **Figure S3A**) owing to the use of different LEDs.

We then measured NMR spectra of fF23AzoF during dichromatic pulsed irradiation using the setup described above. Unexpectedly, these initial results indicated that AzoF adopted a *Z*-enriched PSS instead of an *E*-enriched PSS observed for isolated AzoF. We suspected that incorporated AzoF in aqueous buffer might show a different photophysical behavior than isolated AzoF in DMSO-$d_6$. Thus, we evaluated the switching behavior of $^{15}$N-fF23AzoF by following the absorbance at the maximum of the π→π* transition (333 nm) over time during monochromatic constantly powered irradiation. For this, we put the cuvette containing $^{15}$N-fF23AzoF directly in front of the respective LED output (**Figure S15**). As a result we determined isomerization rates of $k^{365}$ ~ $2.04\times10^{-2}$ s$^{-1}$ (**Figure S16B**) and $k^{524}$ ~ $0.13\times10^{-2}$ s$^{-1}$ (**Figure S16C**) with half-times of $t_{½(365)}$ ~ 33.9 s and $t_{½(524)}$ ~ 517.3 s. Compared to the rates obtained for isolated AzoF (**Figure S14**, $k^{365}$ ~$4.9\times10^{-2}$ s$^{-1}$ and $k^{530}$ ~$1.5\times10^{-2}$ s$^{-1}$), $k^{365}$ and $k^{524}$ are 2–12-fold slower. Moreover, both PSSs are *Z*-enriched with 10*E*:90*Z* for 365 nm irradiation and 41*E*:59*Z* for 530 nm irradiation explaining why a *Z*-enriched PSS was obtained with dichromatic irradiation. Moreover, the initial NMR spectra lacked an additional signal for the third AzoF species. Hence, we suspect that the appearance of the third species is highly dependent on the wavelengths and the intensities of both LEDs used for dichromatic irradiation.

For the time being, we thus discontinued our NMR experiments since we consider it necessary to initially investigate in future studies, which wavelength compositions and which intensities of irradiation are able to manifest the third AzoF species. Additionally, further technical development of the irradiation setup might be necessary to detect the AzoF species in NMR experiments.

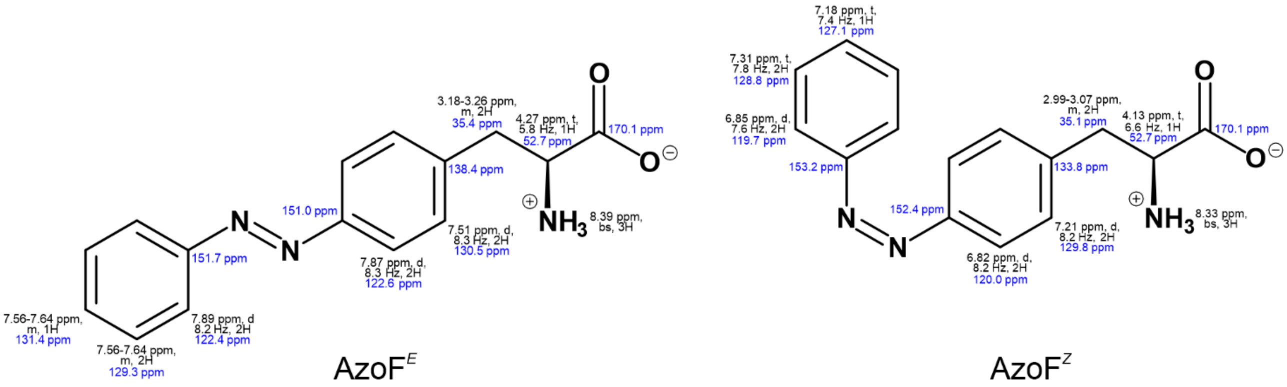


**Figure S9.** Assignment of NMR-signals to the two isomers of AzoF (*E* and *Z*) in 10 mM DMSO-$d_6$ at 25°C. $^1$H chemical shift, multiplicity, coupling constants and integral are highlighted black and $^{13}$C blue.

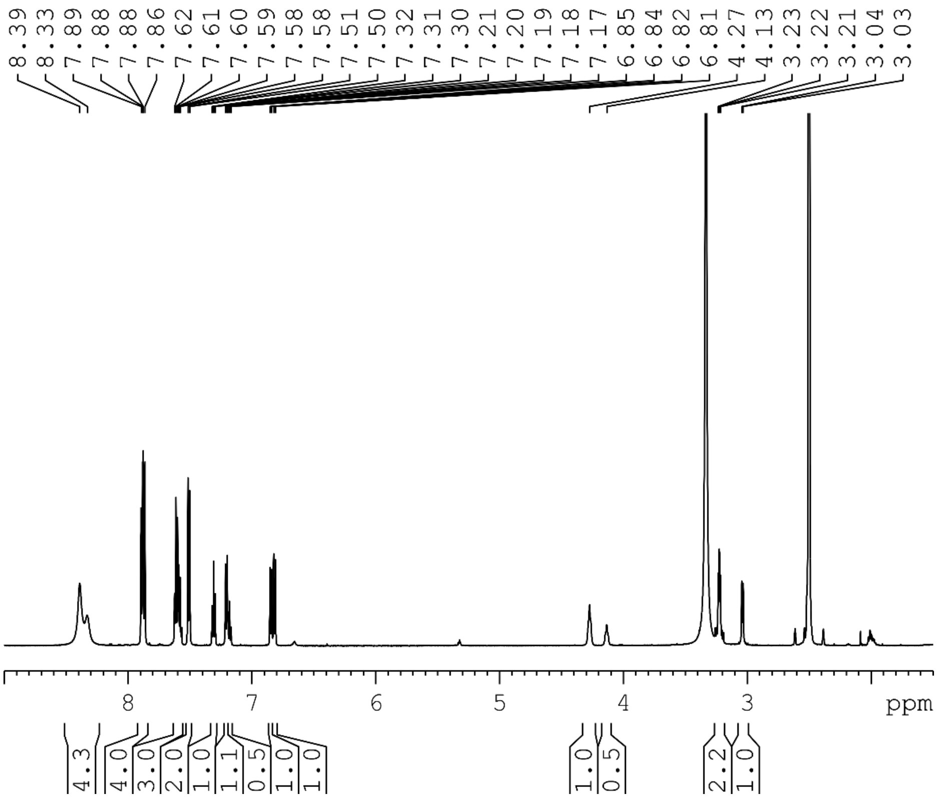

**Figure S10.** 1D-$^1$H spectrum (ns = 32) of 10 mM AzoF in DMSO-$d_6$. The integrals are calibrated on the *E* isomer of AzoF.

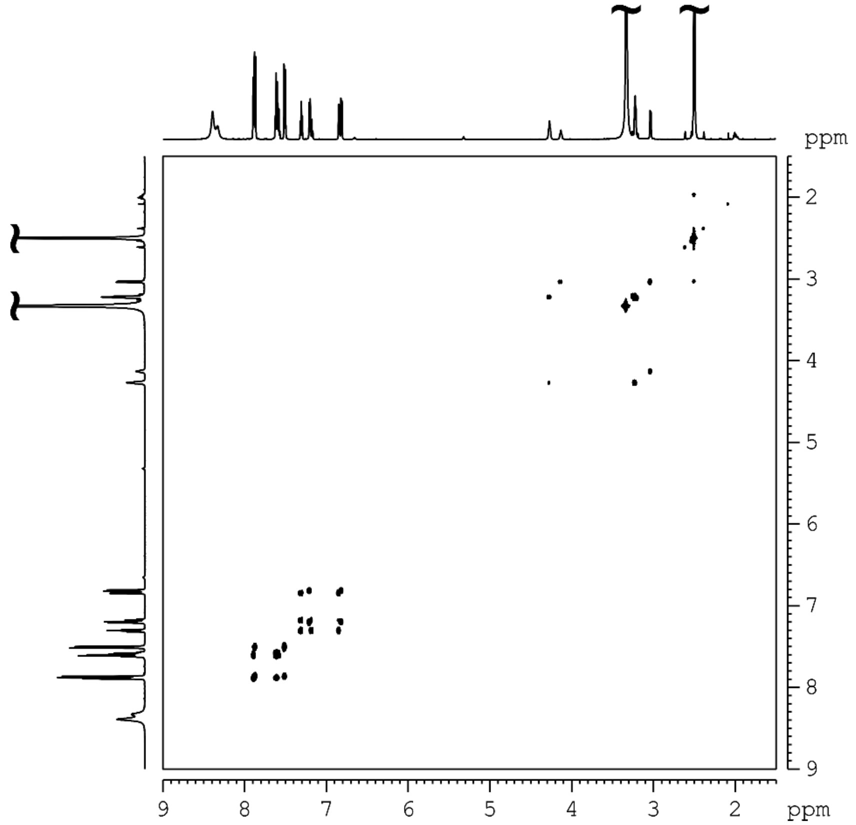


**Figure S11.** 2D-$^{1}$H, $^{1}$H COSY spectrum of 10 mM AzoF in DMSO-$d_6$.

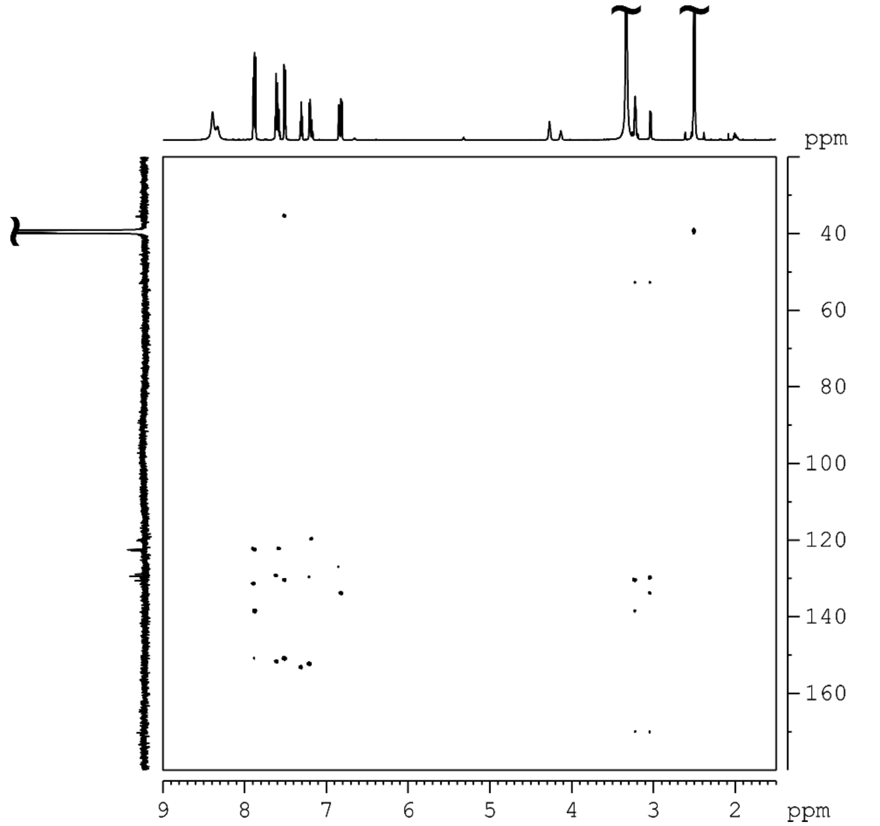


**Figure S12.** 2D-$^{1}$H, $^{13}$C HMBC spectrum of 10 mM AzoF in DMSO-$d_6$.

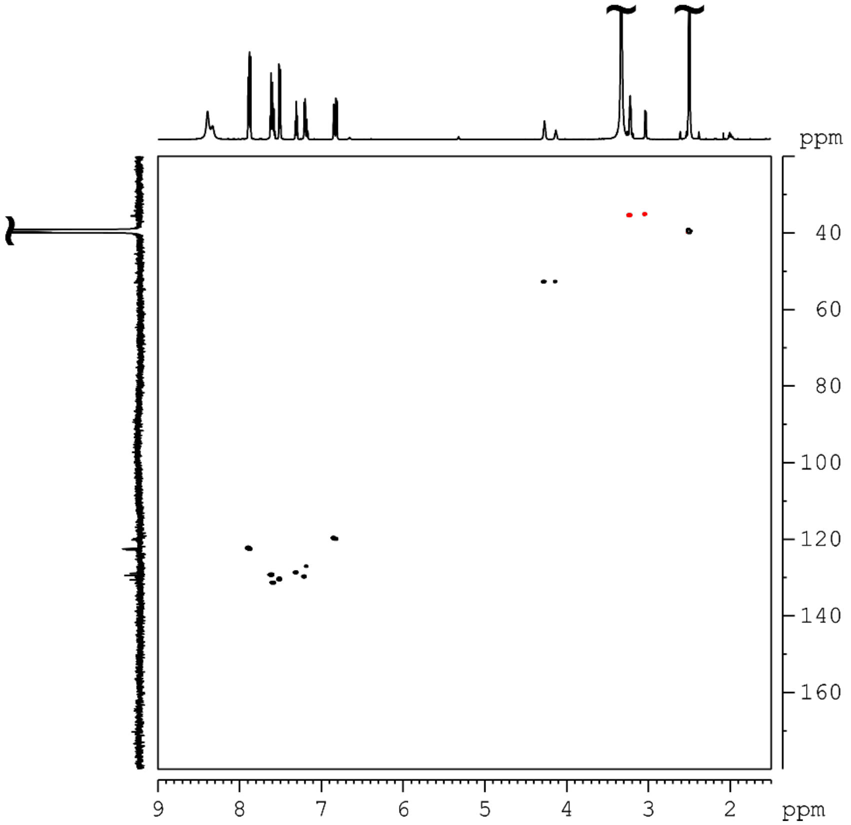


**Figure S13.** 2D-$^{1}H$, $^{13}C$ HSQC spectrum of 10 mM AzoF in DMSO-$d_6$.

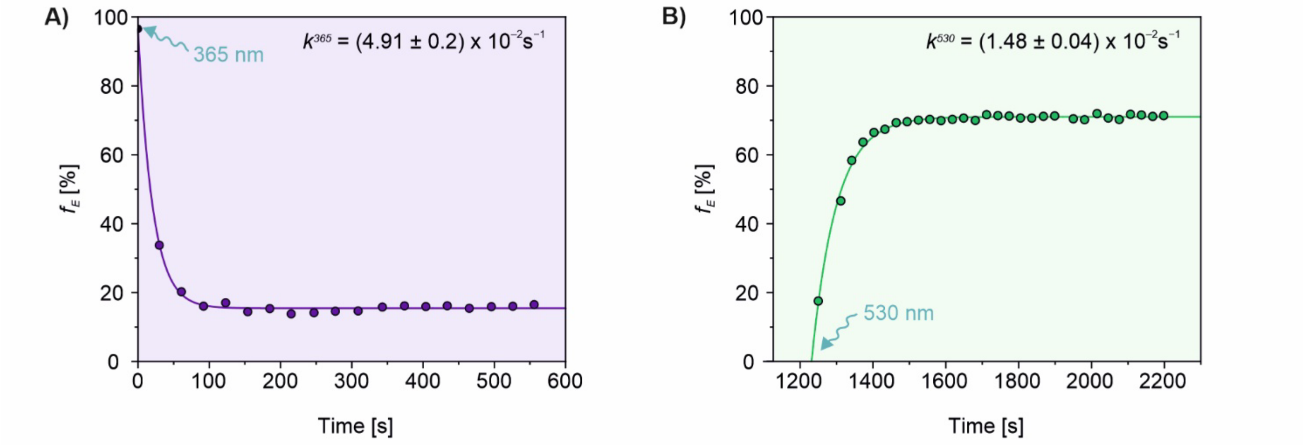


**Figure S14.** Time-resolved 1D-$^{1}H$ NMR spectra (ns = 1) directly following the isomerization of 10 mM AzoF in DMSO-$d_6$ upon monochromatic constantly powered irradiation. A) Excitation of the $\pi\rightarrow\pi^*$ transition with 365 nm initiated a *Z*-enriched PSS. B) Excitation of the $n\rightarrow\pi^*$ transition with 530 nm initiated an *E*-enriched PSS. Mono-exponential fitting obtained the rate constants $k^{365}$ and $k^{530}$ ± S.E..

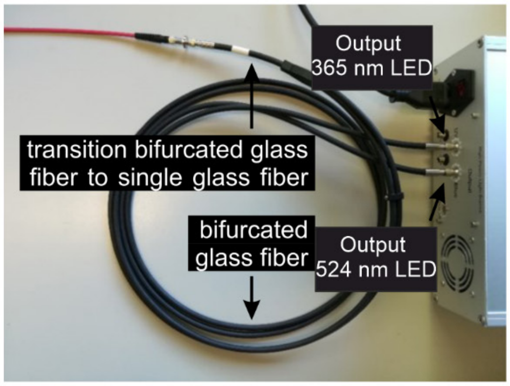

**Figure S15.** Illustration of the NMR irradiation setup that allowed for the monochromatic constantly powered irradiation as well as the dichromatic pulsed irradiation. Samples can be irradiated in an NMR tube containing a single glass fiber that is connected to the LED outputs via a bifurcated glass fiber or in a UV/Vis cuvette placed directly in front of the respective LED output.

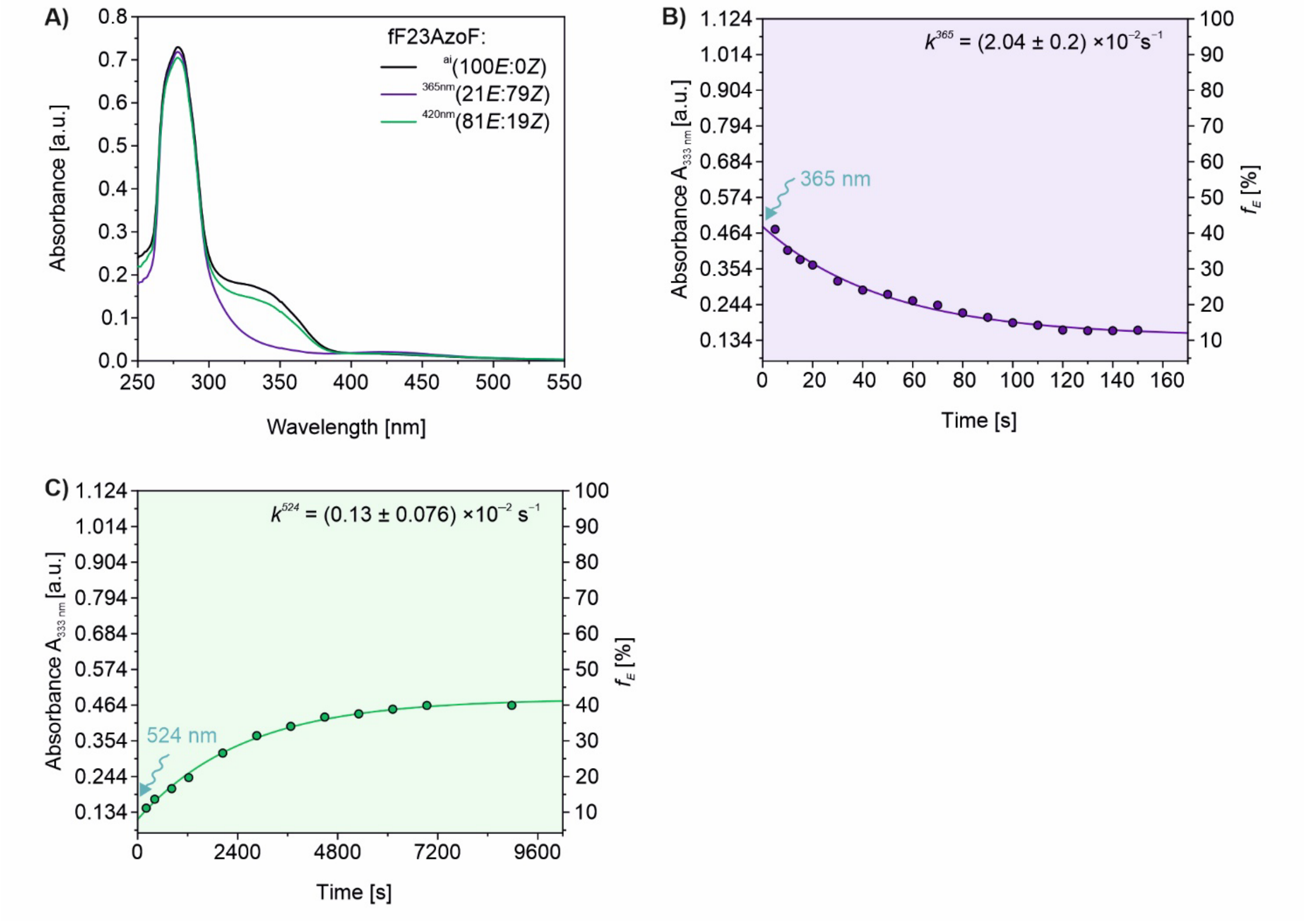


**Figure S16.** Photophysical behavior of 40 µM $^{15}$N-fF23AzoF (in 25 mM NaP pH 7.4, 150 mM NaCl, 25 °C) during monochromatic constantly powered irradiation generated by the NMR irradiation device (**Figure S15**). A) UV/Vis spectra $^{15}$N-fF23AzoF in its as-isolated state (ai, black), in its PSS$^{365}$ (violet), and in its PSS$^{420}$ (green) to confirm incorporation and integrity of AzoF. Note: The here used LEDs are the same as used in ref[1,2]. As a result, the obtained PSSs resemble the previously reported values but differ from those measured in this manuscript, which were obtained using different LEDs (for details see p. 16). C, D) Progress curve following the change of the *E* isomer at 333 nm during 365 nm (C) and 524 nm (D) irradiation directly in front of the LED output of the NMR setup. After certain time-points irradiation was stopped and a UV/Vis spectrum of the sample was recorded. See **Figure S5** for a description of the approximation of $f_E$. Mono-exponential fitting obtained the rate constants $k^{365}$ and $k^{524}$ ± S.E..

#### EXPERIMENTAL SECTION

##### Chemicals and other resources

All reagents and solvents other than listed in **Table S3** were purchased in analytical grade or higher from commercial sources and were used without further purification, if not otherwise stated. *Escherichia coli* BL21 Gold (DE3) cells were further maintained by following the manufacturer's guidelines for the preparation of chemically competent cells.

**Table S3. Key resources**

| Reagent or resource | Source | Identifier |
|---|---|---|
| **Chemicals, Cells, and Recombinant Proteins** | | |
| AzoF | This paper | N/A |
| ProFAR | This Paper | N/A |
| HisG/IE | This paper | N/A |
| HisA | This paper | N/A |
| *E. coli* BL21 Gold (DE3) | Agilent Technologies | # 230132 |
| HisF (wt and fS55AzoF) | This paper | N/A |
| HisH (wt and hW123AzoF) | This paper | N/A |
| **Recombinant DNA** | | |
| pET28a_HisF wt and fS55AzoF | ref [1] | N/A |
| pET28a_HisH wt and hW123AzoF | ref[1,2] | N/A |
| pEVOL_AzoF-RS | Peter Schultz (Scripps Research Institute, La Jolla[3] | N/A |
| **Software and Algorithms** | | |
| Origin 2024 | OriginLab | https://www.originlab.com |
| Pymol 2.0 | Schrödinger 2015 | https://www.pymol.org |
| TopSpin 3.2 & 4.0 | Bruker | https://www.bruker.com |

##### Irradiation setups

Illumination of samples was performed with different irradiation setups (**Table S4**) depending on the experiment. Exact irradiation conditions are given in the respective caption or table heading related to the experiment. All setups were developed in-house.

The UV/Vis setup allowed for the monochromatic and dichromatic constantly powered or pulsed irradiation with 365 nm and 420 nm as illustrated in **Figure S2** and was used for the majority of experiments, unless otherwise stated. This setup contains two output channels that enable simultaneous connection to a 365 nm LED and a 420 nm LED. Adjustable parameters include: current (100-800 mA), frequency (1-10,000 Hz), and duty cycle (10-100%). The LEDs were installed perpendicular to the UV/Vis spectrophotometer measurement beam to allow for simultaneous irradiation and detection of absorbance of the sample. Constantly powered as well as pulsed irradiation was conducted at 365 nm (400 mA) or at 420 nm (400 mA) either at one wavelength or alternating between 365 nm (400 mA) and 420 nm (400 mA) at a set duty cycle ratio of 1:1 and frequencies in the range from 1 Hz to 1000 Hz. Detailed irradiation conditions are listed in the captions and table notes associated with the respective experiment.

The individual setups, as used previously,[1,2] contained either a 365 nm or a 420 nm LED and can be used to irradiate all kinds of samples. In this study, they were used to investigate the integrity and switching behavior of AzoF incorporated into $^{15}$N-fF23AzoF. Only monochromatic constantly powered irradiation was feasible. For irradiation, the sample was placed directly in front of the LED and irradiation with 365 nm (LED Engin Q65113A2058, 20 V 800 mA) or 420 nm (Avonec 1W410420m, 12 V, 350 mA) was maintained for 1–3 min to ensure that the PSS was established.

**Table S4. Irradiation setups used in this work.**

| Irradiation setup | λ [nm] | LED | Mode of irradiation | Applied settings | Optical power output |
|---|---|---|---|---|---|
| UV/Vis setup | 365 | SSC VIOSYS CUN66A1B UV Z5 series | Constant / pulsed | 400 mA | 600 mW[a] |
| | 420 | Intelligent LED Solutions ILH-XC01-S410-SC211-WIR200 | Constant / pulsed | 400 mA | 700 mW[a] |
| Individual setups | 365 | LED Engin Q65113A2058 | Constant | 850 mA | 250 mW cm$^{-2}$ [b] |
| | 420 | Avonec 1W410420m | Constant | 350 mA | 60 mW cm$^{-2}$ [b] |
| NMR Setup | 365 | LG Innotek LEUVA66X00RV00 | Constant | 1 A | 2060 mW[c] |
| NMR Setup | 530 | —[d] | Constant | 1 A | n.d. |
| NMR Setup beta version | 365 | Inolux, IN-C68QA(X)TM UV series, 6868 UV LED | Constant / pulsed | 1 W | 10 W[c] |
| NMR Setup beta version | 524 | LED Engin, LZ4-00G108 | Constant / pulsed | 1 W | 10 W[c] |

[a] Intensity was determined with a High Sensitivity Thermophile sensor PM3 Top Assy (Coherent, Range 500 µW–2 W). [b] Intensity was determined with a RM-12 radiometer and UVA$^{+}$ sensor (Optysec, Range 0–2000 mW cm$^{-2}$). [c] Manufacturer specification. *n.d.* irradiation intensity was not determined. [d] Manufacturer could not be retrieved.

The NMR setup[4] that allowed for the monochromatic constantly powered irradiation with 365 nm and 530 nm was used to determine the isomerization rates of isolated AzoF via NMR measurements. This setup consists of a power supply, a potentiometer to adjust the current through the LED and a transistor controlled by the spectrometer. The LED output is coupled to an optical fiber with a roughened tip that is inserted in a coaxial inlet inside the NMR tube.

A beta version of the original NMR setup was designed in collaboration with HK Testsysteme GMbH and allowed for the monochromatic constantly powered irradiation with 365 nm and 524 nm as well as for the dichromatic pulsed irradiation with both wavelengths (**Figure S15**). In this setup, two LEDs are connected with the illumination insert as described previously[4] using a bifurcated glass fiber (Thorlabs; fiber type: BFY1000HS02, 0.39 NA, 300 - 1200 nm, 1000 µm core) and the corresponding SMA adapters to maximize the light coupling to the NMR sample. Irradiation during NMR spectroscopic measurements was facilitated by use of a single glass fiber connected to the LED outputs via the bifurcated glass fiber that was directly inserted into the NMR tube. Alternatively, a cuvette containing the sample was irradiated directly in front of the LED output. It allows a switching frequency from < 1 Hz to several kHz and a light output between 1 W and 7 W can be adjusted. In this work, monochromatic constantly powered irradiation was performed with a radiant flux of 1 W. This setup was used to determine the isomerization rates of $^{15}$N-fF23AzoF via UV/Vis measurements.

#### Synthesis of AzoF

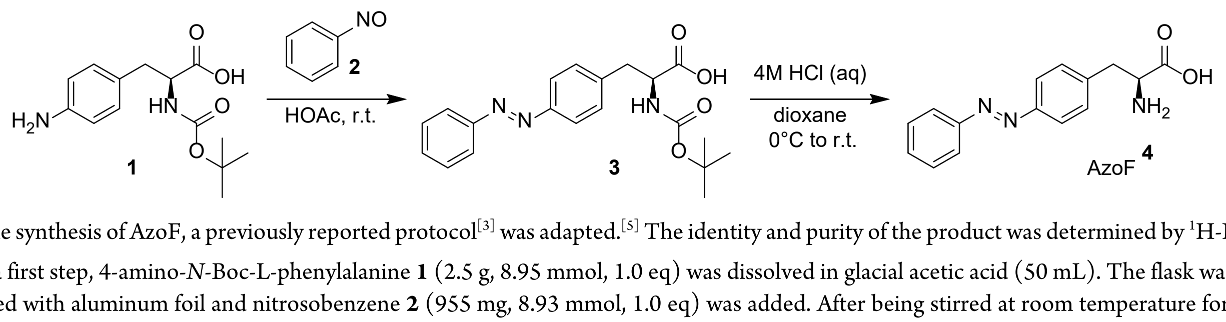


For the synthesis of AzoF, a previously reported protocol[3] was adapted.[5] The identity and purity of the product was determined by $^{1}$H-NMR.

In a first step, 4-amino-*N*-Boc-L-phenylalanine **1** (2.5 g, 8.95 mmol, 1.0 eq) was dissolved in glacial acetic acid (50 mL). The flask was then covered with aluminum foil and nitrosobenzene **2** (955 mg, 8.93 mmol, 1.0 eq) was added. After being stirred at room temperature for 24 h, the reaction mixture was quenched by being poured onto crushed ice/water (500 mL) and extracted with ethyl acetate (3×100 mL). The combined organic layers were washed with water (2×50 mL) and brine (1×50 mL) and were subsequently dried with $Na_2SO_4$. The volatiles were removed in vacuo and the crude product was purified by an automated flash column chromatography (Biotage® Selekt, Biotage) in DCM with a linear gradient of methanol (0–10%) yielding intermediate **3**, which was directly used in the next step.

To obtain AzoF (**4**), intermediate **3** (1.8 g, 6.44 mmol) was dissolved in dioxane (20 mL) at 0 °C. Then, 4 M HCl (6.6 mL) was added, and the mixture was stirred at room temperature for 24 h. Toluene (50 mL) was added to facilitate the removal of 1,4-dioxane, and the solvent was removed in vacuo. The product was triturated with diethyl ether (4×50 mL) and lyophilized overnight, yielding AzoF (overall yield 54–63%).**$^{1}$H-NMR** (300 MHz, Chloroform-*d*): δ = 7.97–7.83 (m, 4H), 7.72–7.55 (m, 3H), 7.54–7.43 (m, 2H), 4.22 (t, J = 6.4 Hz, 1H), 3.25 (t, J = 6.8 Hz, 2H).

##### Synthesis of ProFAR

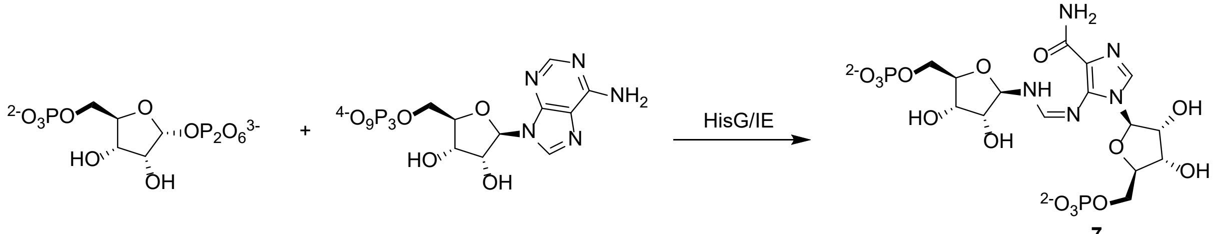


**5** **6** ProFAR **7**

ProFAR was synthesized following a standard protocol developed by Davisson *et al.*[6]. For this, the enzyme HisG/IE was employed to allow for the condensation of 5-phospho-D-ribosyl α-1-pyrophosphate **5** and adenosine triphosphate **6** to yield ProFAR **7** in 50 mM $NH_4COOCH_3$ pH 7.8. The reaction progress was tracked spectrophotometrically[6] and after total conversion, the products were purified with ion exchange chromatography using a POROS HQ 20 column (10 mL, Applied Biosystems) and a gradient of $NH_4COOCH_3$ (50 mM → 1 M). ProFAR concentration was determined at 300 nm ($\varepsilon_{300}$ = 6069 $M^{-1}cm^{-1}$).[7] Highly concentrated and >90% pure ($A_{290}/A_{260}$ = 1.1–1.2 accounts for >95% purity) fractions were lyophilized and stored at −80 °C.

##### Auxiliary enzymes

The auxiliary enzyme HisG/IE was produced by heterologous gene expression in *E. coli* BL21 Gold (DE3) from p_hisGIE_tac as previously described.[6] After transformation, cells were grown in lysogeny broth (LB) medium (2 L) at 37 °C to an $OD_{600}$ of 0.7. Protein expression was induced by addition of 1 mM isopropyl β-D-thiogalactopyranoside (IPTG), and the culture was further incubated at 30 °C overnight. Cells were harvested by centrifugation, resuspended in 50 mM KP pH 7.5, 2.5 mM EDTA, and 1 mM DTT and lysed by sonication. The proteins were subjected to ion exchange chromatography using a MonoQ HR 16/10 column (20 mL, Pharmacia) with a linear gradient of KP (10 → 500 mM). Fractions containing the target proteins with >90% purity as judged by SDS-PAGE analysis were pooled and dialyzed against 50 mM KP pH 7.5, 2.5 mM EDTA, and 1 mM DTT. The protein was dripped into liquid nitrogen for storage at −80 °C.

HisA from *Thermotoga maritima* was produced by heterologous gene expression in *E. coli* BL21 Gold (DE3) from pET21_HisA as previously described.[8] The recombinant protein was purified from the crude extract by heat precipitation (15 min at 73 °C) of host proteins, followed by nickel-affinity chromatography (HisTrap_FF Crude column, 5 mL GE Healthcare) employing a linear gradient of imidazole (1–300 mM) in 50 mM potassium phosphate (KP) pH 7.5, 300 mM NaCl. Fractions containing the pure protein were pooled and dialyzed against 50 mM KP pH 7.5. Based on SDS-PAGE analysis, the purity of all samples was at least 95%. The protein was dripped into liquid nitrogen for storage at −80 °C.

##### Expression and purification of HisF, HisH, hW123AzoF, fS55AzoF, and $^{15}$N-fF23AzoF

Wild type HisH and HisF from *Thermotoga maritima*, as well as associated variants for the alanine scan were recombinantly expressed in *E. coli* BL21 Gold (DE3) and purified following a standard protocol as described previously.[1,2] Briefly, the cells were transformed with the respective plasmids and then grown in an overnight culture of LB medium. The next day, 4 L of LB medium were inoculated with the overnight culture to and $OD_{600}$ of ~0.1, supplemented with 50 mg/L kanamycin and grown at 37 °C. As soon as an $OD_{600}$ of 0.6 was reached, 0.5 mM IPTG was added, and protein expression proceeded at 30 °C overnight. Cells were harvested by centrifugation, resuspended in either 50 mM Tris-HCl pH 7.5, 100 mM NaCl, and 10 mM imidazole (HisF) or 50 mM KP pH 7.5, 100 mM NaCl, and 10 mM imidazole (HisH) and lysed by sonification. *E. coli* proteins were precipitated by a heat step (65 °C, 15 min) and the proteins of interest were obtained from the supernatant after centrifugation. The proteins were subjected to nickel-affinity chromatography (HisTrap_FF Crude column, 5 mL GE Healthcare) and elution was conducted with a linear gradient of imidazole (10-750 mM). Fractions containing the desired proteins were identified by sodium dodecyl sulfate-polyacrylamide gel electrophoresis (SDS-PAGE), pooled and further purified via preparative size-exclusion chromatography (Superdex 75 HiLoad 26/600, GE Healthcare) using 50 mM HEPES pH 7.5, 100 mM NaCl as running buffer. Fractions containing protein with a purity of >90% were identified by means of SDS-PAGE, pooled, concentrated, and dripped into liquid nitrogen for storage at −80 °C.

For the expression of fS55AzoF and hW123AzoF, *E. coli* BL21 Gold (DE3) were co-transformed with the plasmid containing the respective DNA sequence of the target protein and pEVOL_AzoF.[3] The latter harbors the orthogonal aminoacyl-tRNA synthetase adapted for AzoF binding as well as the respective orthogonal tRNA required for AzoF incorporation. After inoculation of 6 L LB medium (supplemented with 30 mg/L chloramphenicol and 50 mg/L kanamycin) with an overnight culture, cells were grown at 37 °C to reach an $OD_{600}$ of 0.6. Centrifugation and subsequent transfer to 600 mL terrific broth (TB) medium allowed for bacterial growth at 37 °C to proceed with higher density. Protein expression was induced at an $OD_{600}$ of ~10 or at least after 6 hours growth by addition of 0.5 mM IPTG, 0.02 % L-arabinose, and 0.8 mM AzoF. The cultures were further incubated at 30 °C overnight. Protein purification was conducted in the dark as described for the wild type enzymes.

For NMR studies, E. coli BL21 Gold (DE3) were co-transformed with pET28a_fF23AzoF and pEVOL_AzoF. Cells were grown at 37 °C in 250 mL LB medium supplemented with 30 mg/L chloramphenicol and 50 mg/L kanamycin to reach an $OD_{600}$ of 0.6—0.8. The culture was then divided into ten 25 mL portions, which were each transferred to 1 L minimal medium supplemented with the respective antibiotics as well as 0.5 g/L $^{15}NH_4Cl$ for 15N-NMR labeling. The cells were further grown at 37 °C to an $OD_{600}$ of 0.7, at which gene expression was induced by addition of 0.5 mM IPTG, 0.02 % L-arabinose and 0.8 mM AzoF. The cultures were further incubated at 30 °C overnight. The following steps were conducted in the dark. The cells were harvested by centrifugation, resuspended in 50 mM sodium phosphate (NaP) pH 7.4, 400 mM

NaCl, and 10 mM imidazole and lysed by sonification. $^{15}$N-fF23AzoF was purified using a heat step (65 °C, 10 min) followed by a nickel-affinity chromatography (HisTrap_FF Crude column, 5 mL GE Healthcare). Elution was conducted with a linear gradient of imidazole (10-750 mM) in 25 mM NaP pH 7.4, 150 mM NaCl. Fractions containing the desired protein were pooled and subjected to digestion with TEV protease in 25 mM NaP pH 7.4, 150 mM NaCl, and 1 mM DTT. Subsequently, nickel-affinity chromatography was used to remove the cleaved $His_6$-tag. $^{15}$N-fF23AzoF was yielded from the flow-through, concentrated, dripped into liquid nitrogen and stored at −80 °C.

#### UV/Vis analysis

UV/Vis spectra of hW123AzoF and fS55AzoF were recorded in an Agilent 8453 spectrophotometer in the range of 190–1100 nm. All measurements were performed with 10–20 µM protein in 50 mM HEPES (pH 7.5), 100 mM NaCl in a 1cm quartz cuvette at room temperature unless otherwise stated. Samples were either pre-irradiated for at least 1 min to ensure that the PSS was reached and measured in the dark or they were irradiated during the measurement using the UV/Vis setup (pp. 16; **Table S4**; **Figure S2**). Detailed irradiation conditions are listed in the figure captions and table notes associated with the respective experiment, such as cycle times for time-resolved spectra.

To ensure the introduction and integrity of AzoF in $^{15}$N-fF23AzoF, spectra of the as-isolated protein as well as after exposure to monochromatic constantly powered irradiation at 365 nm and 420 nm were measured. Spectra were recorded of 40 µM protein (in 25 mM NaP pH 7.4 and 150 mM NaCl) at room temperature in the range of 250–600 nm using a 1 cm quartz cuvette and a JASCO V650 spectrophotometer. Irradiation took place by removing the cuvette from the photometer and placing it directly in front of the individual irradiation setups (**Table S4**). To determine the rate constants of AzoF isomerization in $^{15}$N-fF23AzoF, the protein was instead irradiated with the NMR setup beta version (**Table S4**; **Figure S15**). For this, the 1 cm quartz cuvette containing the sample was placed directly in front of the LED output of the NMR setup. At certain time-points, irradiation was stopped, and a UV/Vis spectrum of the sample was obtained in the JASCO V650 spectrophotometer.

All spectra were baseline corrected at 600 nm. To determine the rates of isomerization *k* the absorbance values at the $\pi \rightarrow \pi^*$ transition (322 nm) were plotted against time. Exponential decay of the absorbance was then fitted with **Equation S1**

$$y = y_{\mathrm{i}} + A\, e^{-\mathrm{kt}}$$

**Equation S1**

in which *t* is the time, $y_i$ is the *y* value at infinite times (also named plateau), *A* is the span of the exponential curve between $y_i$ and $y_0$ (the y value when time = 0), and *k* is the rate constant. Exponential growth of the absorbance was fitted with **Equation S2.**

$$y = y_0 + A\,(1 - e^{-\mathrm{kt}})$$

**Equation S2**

The half-life of isomerization $t_{1/2}$ was then derived from the isomerization rate *k* with **Equation S3**.

$$t_{1/2} = \frac{\ln(2)}{k}$$

**Equation S3**

#### Estimation of the *E*:*Z* ratio from UV/Vis spectra

The *E*:*Z* ratios of hW123AzoF, fS55AzoF and $^{15}$N-fF23AzoF were estimated following an expanded procedure based on previously published protocols[9] and is shown exemplarily in **Figure S6**. In the first step, several UV/Vis spectra spanning various time-points between the as-isolated state and a PSS were selected from one experiment and deconvoluted with the multiple peak fit tool in Origin 2024 using the Gaussian function in **Equation S4**

$$y = A\, e^{-\frac{(x-x_c)^2}{2w^2}}$$

**Equation S4**

in which *y* is the measured absorbance, *A* is the amplitude of the Gauss curve, *x* is the wavelength, $x_c$ is the wavelength at the center of the Gauss curve, and *w* is the width of the Gauss curve at $\frac{A}{2}$. We thereby focused on the $\pi \rightarrow \pi^*$ and $n \rightarrow \pi^*$ peaks and, hence, performed the deconvolution between 310 nm and 600 nm. As a result, a cumulative Gauss fit of both peaks was obtained simulating the respective UV/Vis spectrum.

In the second step, we used the cumulative Gauss fits to obtain a first estimate of the *E*:*Z* ratios. For this, we assumed that the as-isolated protein comprises 100% of the *E* isomer as AzoF should be thermally equilibrated and used the spectrum of the as-isolated state as reference. To determine the fraction of *E* in the other selected UV/Vis spectra at various time-points, the spectrum of the 100% *Z* isomer was simulated with the simple curve math tool in Origin 2024 using **Equation S5**

$$A_Z = \frac{A_i - A_E x\, f_E}{1 - f_E}$$

**Equation S5**

in which $A_Z$ is the simulated spectrum of the 100% *Z* isomer, $A_E$ is the cumulative Gauss fit of the 100% *E* reference spectrum, $A_i$ is the cumulative Gauss fit of a spectrum at a certain time point, and $f_E$ is the fraction of the *E* isomer. The fraction of $f_E$ was obtained in this simulation by manual adjustment until the signal of the $\pi \rightarrow \pi^*$ absorbance band approximated zero.

In the final step, we used the estimated $f_E$ values to produce a standard curve for each experiment. By this, we intended to improve the quality of the estimation and simultaneously simplify the process for all measured spectra within an experiment. To this end, the $f_E$ values were plotted against the absorbance value at 322 nm for each original UV/Vis spectrum that was selected in the first step. Linear regression of this standard curve then allowed us to estimate $f_E$ of any spectrum within the experiment using the linear fit equation and the absorbance at 322 nm.

#### Determination of ImGPS activity in the dark and during irradiation

The ImGPS reaction was measured by following the turnover of PRFAR to AICAR and ImGP continuously at 300 nm [$\Delta\varepsilon_{300}$(PRFAR-AICAR) = 5637 $M^{-1}cm^{-1}$] with an Agilent 8453 Spectrophotometer. Reaction conditions included 0.6 µM HisA (to turn over ProFAR to PRFAR), approximately 40 µM (saturated) ProFAR, and 10 mM (saturated) glutamine in 50 mM Tris-acetate (pH 7.5) at room temperature or at 20 °C. For complex formation, the respective HisF and HisH monomers were mixed at equimolar concentrations in 50 mM Tris-HCl (pH 7.5) and the reaction was started with either 0.1 µM ImGPS(hW123AzoF), 0.2 µM ImGPS(fS55AzoF), or 0.025 µM wt-ImGPS. PRFAR turnover was followed in the dark for approximately 5 min before irradiation of the sample started using the UV/Vis setup (**Table S4; Figure S2**). This allowed for the irradiation either with a constant current at 365 nm, 420 nm or both, or with pulse width modulation (PWM) at 365 nm, 420 nm or both. Irradiation was either maintained until the end of the experiment or was stopped after a certain amount of time and PRFAR turnover was followed in the dark until the reaction was completed. Detailed irradiation conditions including pulse frequencies for PWM measurements are given in the captions and table notes associated with the respective experiment.

Since both substrates were provided in saturation, the reaction occurred with pseudo-zero order kinetics. This means the progress curves followed an almost entirely linear behavior throughout the course of the reaction. Upon the start of irradiation, the reaction velocity was changed but the linear behavior was maintained (cf. **Figures 2–3**). Hence, the reaction velocity *v* of each reaction segment (prior to, during and after irradiation) was determined using a linear fit over the entire segment. Multiple biological and technical replicates were summarized by calculating the mean and the standard error of mean (S.E.M.) of *v*.

The light regulation factor (LRF) was determined for each individual experiment using **Equation S7**

$$LRF = \frac{v_1}{v_2} \; (v_1 > v_2)$$

**Equation S7**

in which $v_1$ is assigned to the reaction segment with the faster reaction velocity, and $v_2$ is assigned to the reaction segment with the slower reaction velocity. The LRF of multiple biological and technical replicates was derived as the mean ± S.E.M. of the individual LRFs.

#### Nuclear magnetic resonance (NMR) spectroscopy of AzoF

Detailed NMR spectroscopic investigations were performed on a Bruker Avance III HD 600 (600.03 MHz) with a 5 mm fluorine selective TBIF probe or on a Bruker Avance NEO 600 (600.03 MHz) with a 5 mm CryoProbe Prodigy BBO. For time-resolved NMR spectroscopic investigations with irradiation the reported setup with 5 mm amberized thin wall NMR tubes was used.[4]

For sample preparation, an oven dried 5 mm amberized thin wall NMR tube was charged with the appropriate amount of AzoF (10 mM) and the DMSO-$d_6$ (0.3 mL) was added *via* a syringe. Before NMR measurement, AzoF was exposed to monochromatic constantly powered irradiation at 25 °C using the glass fiber of the NMR setup or the NMR setup beta version (**Table S4; Figure S15**) that was directly inserted into the NMR tube. The change of the *E* isomer fraction was determined from the respective signals in the 1D-$^1$H NMR spectra (ns = 1) during 368 nm and 524 nm irradiation. All spectra were referenced to the solvent residual signal: DMSO-$d_6$: $\delta(^1H) = 2.50$ ppm. NMR data was processed, evaluated, and plotted with TopSpin 3.2 and 4.0 software. Further plotting of the spectra was performed with Origin 2024 and Corel Draw 2020 software.

#### REFERENCES


[1] A. C. Kneuttinger, K. Straub, P. Bittner, N. A. Simeth, A. Bruckmann, F. Busch, C. Rajendran, E. Hupfeld, V. H. Wysocki, D. Horinek et al., *Cell Chem. Biol.* **2019**, *26*, 1501–1514.

[2] A. C. Kneuttinger, C. Rajendran, N. A. Simeth, A. Bruckmann, B. König, R. Sterner, *Biochemistry* **2020**, *59*, 2729.

[3] M. Bose, D. Groff, J. Xie, E. Brustad, P. G. Schultz, *J. Am. Chem. Soc.* **2006**, *128*, 388.

[4] C. Feldmeier, H. Bartling, E. Riedle, R. M. Gschwind, *J. Magn. Reson.* **2013**, *232*, 39.

[5] C. Hiefinger, S. Mandl, M. Wieland, A. Kneuttinger in *Methods in Enzymology* (Ed.: A. K. Shukla), Academic Press, **2023**, pp. 247–288.

[6] V. J. Davisson, I. L. Deras, S. E. Hamilton, L. L. Moore, *J. Org. Chem.* **1994**, *59*, 137.

[7] T. J. Klem, V. J. Davisson, *Biochemistry* **1993**, *32*, 5177.

[8] F. List, M. C. Vega, A. Razeto, M. C. Häger, R. Sterner, M. Wilmanns, *Chem. Biol.* **2012**, *19*, 1589.

[9] a) J. Calbo, C. E. Weston, A. J. P. White, H. S. Rzepa, J. Contreras-García, M. J. Fuchter, *J. Am. Chem. Soc.* **2017**, *139*, 1261; b) K. Rustler, P. Nitschke, S. Zahnbrecher, J. Zach, S. Crespi, B. König, *J. Org. Chem.* **2020**, *85*, 4079.

[10] J. Luo, S. Samanta, M. Convertino, N. V. Dokholyan, A. Deiters, *ChemBioChem* **2018**, *19*, 2178.

[11] W. L. Butler, S. B. Hendricks, H. W. Siegelman, *Photochem. Photobiol.* **1964**, *3*, 521.

[12] a) J. P. Wurm, S. Sung, A. C. Kneuttinger, E. Hupfeld, R. Sterner, M. Wilmanns, R. Sprangers, *Nat. Commun.* **2021**, *12*, 2748; b) J. M. Lipchock, J. Patrick Loria, *Biomol. NMR Assign.* **2008**, *2*, 219; c) J. Lipchock, J. P. Loria, *J. Biomol. NMR* **2009**, *45*, 73.
